# A Single Intranasal Dose of Bacterial Therapeutics to Calves Confers Longitudinal Modulation of the Nasopharyngeal Microbiota

**DOI:** 10.1101/2022.01.03.474870

**Authors:** Samat Amat, Edouard Timsit, Matthew Workentine, Timothy Schwinghamer, Frank van der Meer, Yongmei Guo, Trevor Alexander

## Abstract

To address the emergence of antimicrobial-resistant pathogens in livestock, microbiome-based strategies are increasingly being sought to reduce antimicrobial use. Here, we describe the intranasal application of bacterial therapeutics (BTs) for mitigating bovine respiratory disease (BRD) and used structural equation modeling to investigate the causal networks after BT application. Beef cattle received i) an intranasal cocktail of previously characterized BT strains, ii) an injection of metaphylactic antimicrobial (tulathromycin), or iii) intranasal saline. Despite being transient colonizers, inoculated BT strains induced longitudinal modulation of the nasopharyngeal bacterial microbiota while showing no adverse effect on animal health. The BT-mediated changes in bacteria included reduced diversity and richness and strengthened cooperative and competitive interactions. In contrast, tulathromycin increased bacterial diversity and antibiotic resistance, and disrupted bacterial interactions. Overall, a single intranasal dose of BTs can modulate the bovine respiratory microbiota, highlighting that microbiome-based strategies have the potential in being utilized to mitigate BRD in feedlot cattle.

## Introduction

Bovine respiratory disease (BRD) is the most significant health condition affecting beef calves, and accounts for economic losses due to costs associated with treatment and prevention, and reduced productivity (1, 2). In North America, the management of beef cattle typically involves their shipment from pastures to feedlots for production. During feedlot placement, cattle are most susceptible to BRD, with the majority of cases occurring within the first 60 days of feedlot placement (3). Primary viral infections or stressors, that include weaning, shipping, and co-mingling with new pen mates, are proposed to reduce host immunity during transition to feedlots (4). Consequently, opportunistic bacterial pathogens residing in the upper respiratory tract proliferate and translocate to the lungs causing bronchopneumonia (5). The main BRD-associated bacteria include *Mannheimia haemolytica*, *Pasteurella multocida*, *Histphilus somni*, and *Mycoplasma bovis* (6). As a result of increased BRD susceptibility, commercial feedlots rely on antimicrobial-driven approaches to prevent BRD infections in cattle (7).

Long-acting injectable antimicrobials are commonly administered to cattle entering feedlots (i.e. metaphylaxis) for BRD prevention (8). For example, the macrolide tulathromycin was used at feedlot entry by 45.3% of feedlots in the United States for BRD mitigation (9). Metaphylactic antimicrobials treat lung infections that may be prevalent in calves entering feedlots and also prevent infection during the course of their bioactivity. It is also likely that metaphylactic antimicrobials reduce prevalence and proliferation of BRD pathogens in the upper respiratory tract, a prerequisite to lung translocation (10). However, antimicrobial resistance in BRD pathogens has increased over the last ten years (11–13) with resistance to tulathromycin recently being detected in more than 70% of *M. haemolytica* and *P. multocida* isolated from feedlot calves (14). In addition, resistance elements have been detected in mobile elements from the BRD-associated *Pasteurellaceae* family, conferring multi-drug resistance to antimicrobials used for both prevention and treatment of BRD in feedlot cattle (15, 16). Antimicrobial resistance in BRD pathogens therefore threatens the efficacy of currently used antimicrobials in beef cattle. Alternatives to antimicrobials are therefore needed for use in novel feedlot management strategies. The respiratory microbiota contributes to host health by providing colonization resistance against pathogens and maintaining homeostasis (17). It is proposed that disruption of the bovine respiratory microbiota can promote the proliferation of BRD pathogens (18). Indeed, several management factors including transportation to a feedlot (19), diet composition (20), and antimicrobial administration (21) alter the upper respiratory tract microbiota of cattle. Recently, we observed that several lactic acid-producing bacteria (LAB) were inversely correlated with *Pasteurellaceae* in the nasopharynx of cattle transported to an auction market and subsequently a feedlot (22). Genera within the LAB order *Lactobacillales* have been shown to be reduced in cattle that develop BRD (23, 24). Additionally, isolates of LAB originating from the bovine respiratory tract have been shown to directly inhibit BRD pathogens (25, 26). These data support that LAB are important community members of the bovine respiratory tract and may be integral to providing colonization resistance against BRD pathogens. Consequently, we have previously developed BTs comprised of six *Lactobacillus* strains that were characterized for their *in vitro* inhibition and exclusion of *M. haemolytica*, and adherence to and immunomodulation of bovine turbinate cells (26). The intranasal inoculation of these BT strains was able to inhibit colonization by *M. haemolytica* in experimentally challenged dairy calves (27). In the present study, we further evaluated the longitudinal effects of these intranasal BTs on the nasopharyngeal microbiota of beef calves after a single intranasal dose. The effects of the BTs were also compared to those of tulathromycin, a common antimicrobial used for metaphylaxis, administered subcutaneously.

## Results

### Calf health and weight gain

Calves were monitored daily for clinical signs of BRD throughout the study. Elevated rectal temperature (≥ 39.7 °C) was detected in five calves (three from CTRL and two from BT; Supplementary Table S1) that recovered in response to a single injection of the antibiotic micotil (tilmicosin). The remainder of the experimental calves were healthy during the course of study. The calves were weighed first at arrival (d-1) and subsequently on a bi-weekly basis for the 42 days of study. The average daily gain was not different among treatments (*P* = 0.506) (Supplementary Fig. S1).

### Prevalence of BRD-associated pathogens determined by NP swab culturing

Presence of *M. haemolytica*, *P. multocida*, and *H. somni* was evaluated by culturing the NP swabs collected during the first 28 days of study (Fig. 1). Overall, *M. haemolytica* had the highest prevalence of the three pathogens on d-1 (24 - 40% across treatments). No significant difference was detected among treatment groups (*P* > 0.05) at any sampling time point for *M. haemolytica*, however, there was a tendency (*P* = 0.06) for reduced prevalence in metaphylaxis (MP)-treated calves on d 7. For *P. multocida*, prevalence in CTRL (10-40%) and BT (5-36%) calves was similar (*P* > 0.05). In MP calves, prevalence of *P. multocida* was lower (range 0-5%) on days 7 and 14 compared to CTRL and BT calves (*P* < 0.05). Overall, the prevalence of *H. somni* remained low throughout the study for all treatment groups and was not different between treatment groups at any sampling point (*P* > 0.05). Only the MP group had colonization rates of 0% for *P. multocida* (days 1, 4, and 14) and *M. haemolytica* (d 1), which occurred after metaphylactic treatment.

**Fig. 1.**
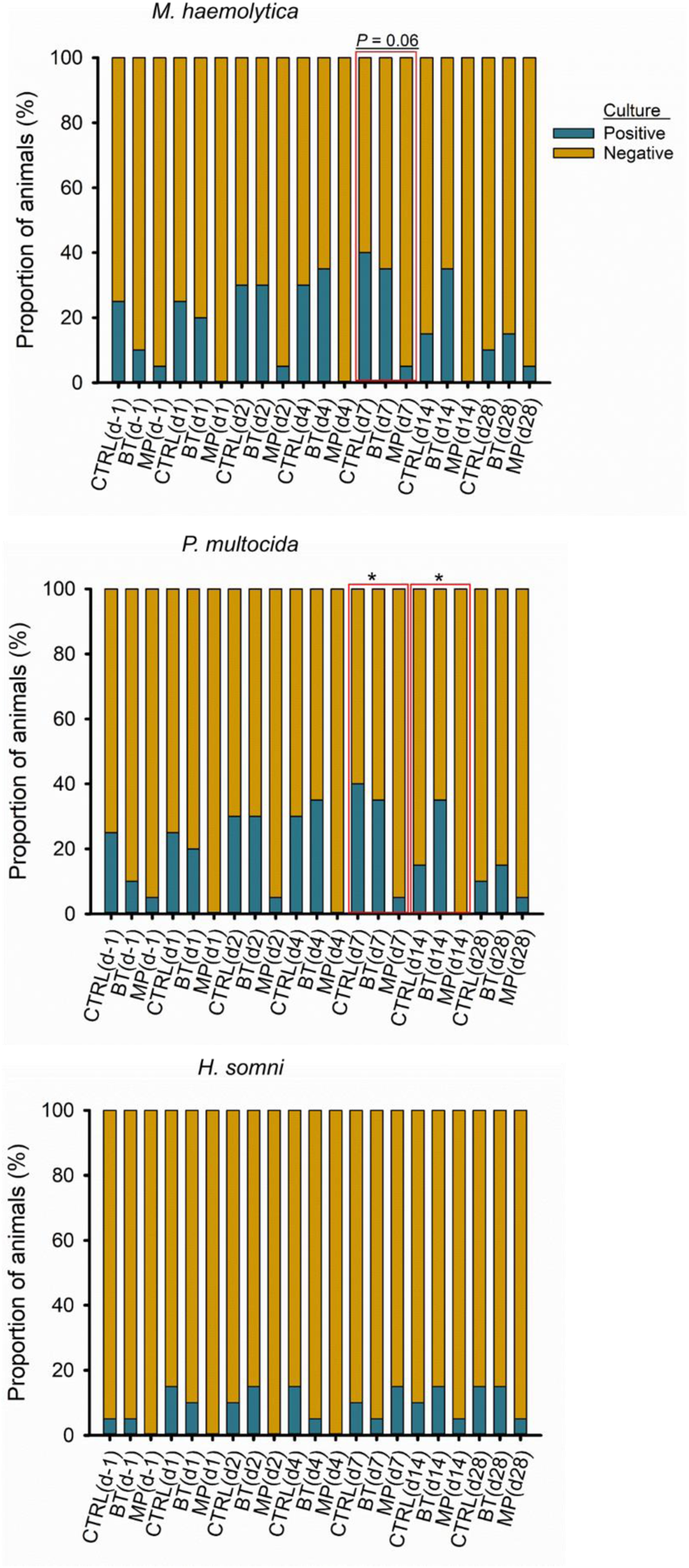
Prevalence of the BRD-associated pathogens in nasopharynx of cattle over the course of 28 days, determined by culturing nasopharyngeal swabs. On day 1, cattle were treated with intranasal bacterial therapeutics (BT), intranasal PBS (CTRL), or subcutaneous tulathromycin (MP) (n = 20 per group). *Significant difference between treatments (P < 0.05).

### Total bacteria and *Lactobacillus* in NP swabs determined by qPCR

The abundances of total bacteria and *Lactobacillus* in nasal swabs was estimated by quantifying the gene copy numbers of general and *Lactobacillus*-specific 16S rRNA (Fig. 2). An interaction between treatment × time affected the total bacterial number (*P* = 0.02; Fig. 2A). Compared to BT and CTRL calves, tulathromycin injection reduced bacteria in NP swabs on days 4, 7, and 42 (*P* < 0.05). Total bacteria in NP swabs of BT and CTRL calves were not different over the course of the study (*P* > 0.05).

**Fig. 2.**
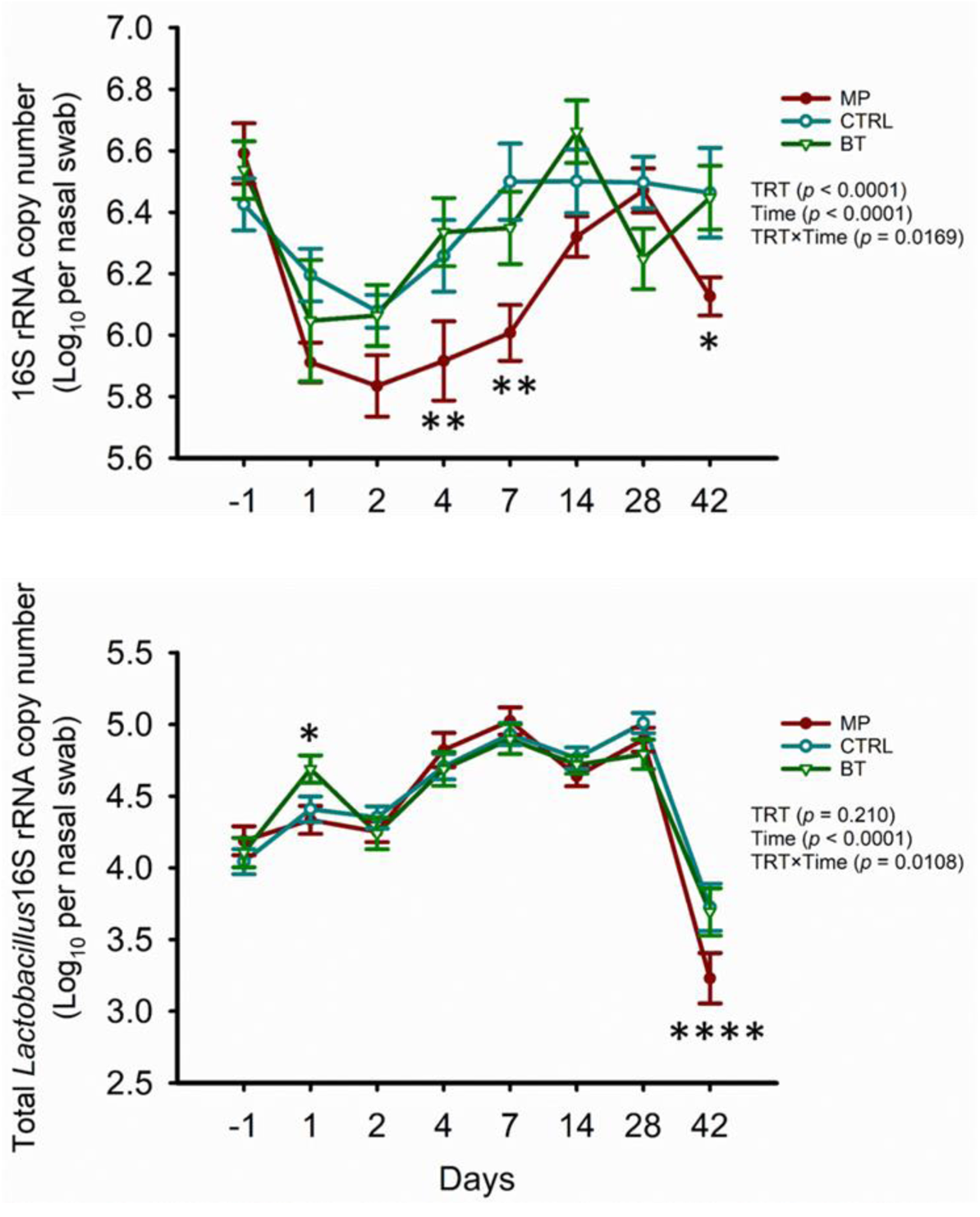
Estimated abundances of total bacteria (top), and *Lactobacillus* (bottom) in nasopharyngeal swab samples obtained from calves that received either intranasal bacterial therapeutics (BT), intranasal PBS (CTRL), or subcutaneous tulathromycin (MP) (n = 20 per group). Gene copy numbers were quantified by qPCR. The results are presented as estimated mean ± SEM. *Significant difference between treatments (* represents *P* < 0.05; ** represents *P* < 0.01; **** represents *P* < 0.0001).

The estimated number of *Lactobacillus* in NP swabs was affected by the interaction of treatment × time (*P* = 0.01). The mean total *Lactobacillus* 16S rRNA copy number per swab from the BT group increased (*P* < 0.01) from d −1 to d 1 (24 h post-BT inoculation) (Fig. 2B), and then decreased to similar levels in CTRL and MP calves by d 2 (*P* > 0.05). On d 42, *Lactobacillus* was reduced in the MP group compared to BT and CTRL calves (*P* < 0.0001).

### Structure and composition of the NP microbiota

#### 16S rRNA gene sequencing overview

The raw SV table contained 10,400 SVs with a total of 7,161,751 reads assigned to 476 samples. The median number of sequences per sample was 15,037 ± 3,482.21 with a minimum of 0 and maximum of 26,536. After filtering, the SV table contained 531 SVs with a total of 6,349,998 reads. The median number of sequences per sample was 13,613 ± 4,100.6 with a minimum of 22 and maximum of 26,027.

#### The community structure of the NP microbiota

PERMANOVA revealed that an interaction of treatment × time had an effect (*R*^2^ = 0.04, *P* = 0.001) on the microbial structure of the NP microbiota (Fig. 3). However, time had a larger effect on microbial structure (*R*^2^ = 0.141, *P* = 0.001) compared to treatment (*R*^2^ = 0.034, *P* = 0.001). As indicated by the DCA plots (Fig. 3) the microbiota tended to cluster by early (d −1 to 2), mid (d 4-7) and late (d14-28) time points. Clustering according to treatment was most evident on d 28 and 42.

**Fig. 3.**
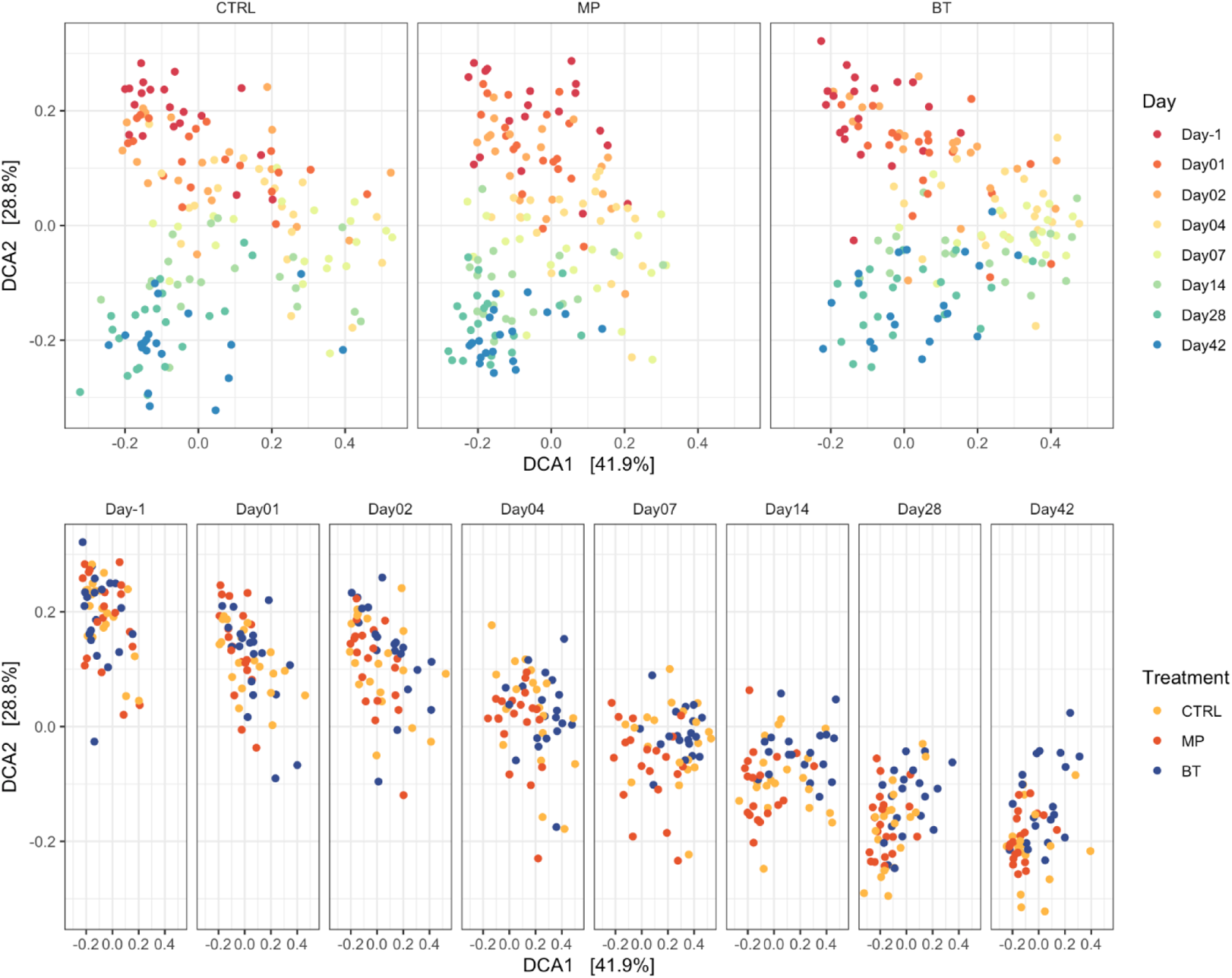
Detrended correspondence analysis (DCA) plots of the Bray-Curtis metric for bacteria in nasopharyngeal samples collected from cattle and visualized by time or treatment. On day 1, cattle were treated with intranasal bacterial therapeutics (BT), intranasal PBS (CTRL), or subcutaneous tulathromycin (MP) (n = 20 per group). The percentages of variation explained by the DCA are indicated on the axes.

Alpha-diversity, as assessed by richness and Shannon diversity index, revealed that both indices were affected by a treatment × time interaction (*P* < 0.05) (Fig. 4). After BT inoculation, the NP microbiota of BT calves had reduced richness throughout the study, compared to CTRL and MP calves (*P* ≤ 0.014), except for on d 7, when richness was similar in BT and CTRL groups (*P* = 0.213). In contrast, MP calves had increased richness on days 7-42 when compared to BT calves and increased richness on days 7 and 14 when compared to CTRL calves (*P* ≤ 0.001). Similar to richness, the Shannon diversity index was reduced in BT calves from days 7-42, compared to CTRL and MP treatments (*P* ≤ 0.0001). The Shannon diversity was greater in MP calves compared to BT and CTRL calves, but only on days 1 and 7 (*P* ≤ 0.016).

**Fig. 4.**
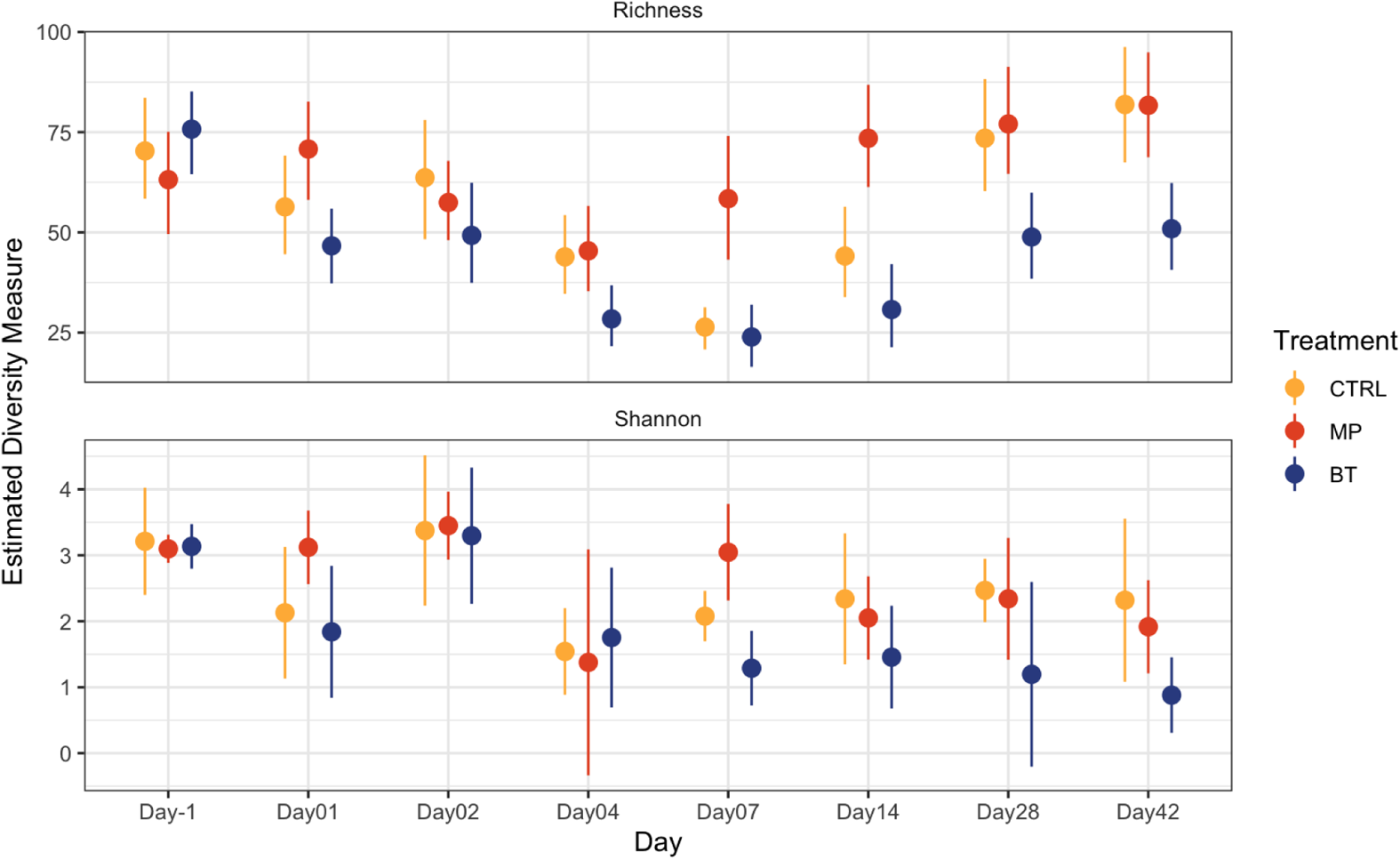
Alpha diversity of bacteria in nasopharyngeal samples collected from cattle over 43 days. On day 1, cattle were treated with intranasal bacterial therapeutics (BT), intranasal PBS (CTRL), or subcutaneous tulathromycin (MP) (n = 20 per group). Top plot shows the mean richness estimate, and the bottom panel shows the mean estimated Shannon diversity. Error bars indicate standard error of the mean.

#### Composition of the NP microbiota

Across time and treatment groups, a total of 14 different bacterial phyla were identified, among which *Proteobacteria* (36.4 %) *Tenericutes* (22.2 %), *Firmicutes* (17.4 %), *Actinobacteria* (12.9 %), and *Bacteroidetes* (9.9 %) were the most relatively abundant, and together constituted 98.8% of the sequences. The diversity of genera within each phylum varied with the relative abundance of a single genus ranging from <1% to 100% of a phylum. Overall, the ten most relatively abundant genera across treatments and time included *Mycoplasma* (22.2%), *Moraxella* (18.8%), *Pasteurella* (3.7%), *Mannheimia* (3.6%), *Corynebacterium_1* (2.7%), *Ruminococcaceae_UCG-005* (2.4%), *Psychrobacter* (2.3%), *Jeotgalicoccus* (2.1%), *Histophilus* (1.1%) and *Planococcus* (1.1%) (Data not shown).

### Changes in microbial composition following BT and tulathromycin treatment

#### Changes in the five most relatively abundant phyla

Noticeable changes in NP microbial composition at phylum level were observed in response to treatment and time effects (Fig. 5). The relative abundance of *Proteobacteria* in CTRL calves varied over the 42 days of study, with a gradual increase in the first 7 days followed by a decline in the remaining 5 weeks of the study. In BT calves, the relative abundance of *Proteobacteria* was similar to CTRL calves, except for on d 28, when it was increased (*P* = 0.02). In MP calves, however, *Proteobacteria* had a lower relative abundance compared to CTRL calves on days 4, 7 and 14 (*P* ≤ 0.008). The relative abundance of *Firmicutus* did not differ between BT and CTRL groups at any sampling time (*P* > 0.05). However, *Firmicutes* was increased in MP calves within the first 14 days of antibiotic injection compared to CTRL calves (*P* < 0.05). Similarly, *Bacteroidetes* became significantly enriched in MP calves on days 7 and 14 relative to the CTRL group (*P* ≤ 0.022). Compared to CTRL calves, the abundance of *Actinobacteia* was reduced in BT calves on d 42 (*P* = 0.011), while it was increased in MP calves on d 1 (*P* = 0.038). The relative abundance of *Tenericutes* was not affected by treatment (*P* > 0.05).

**Fig. 5.**
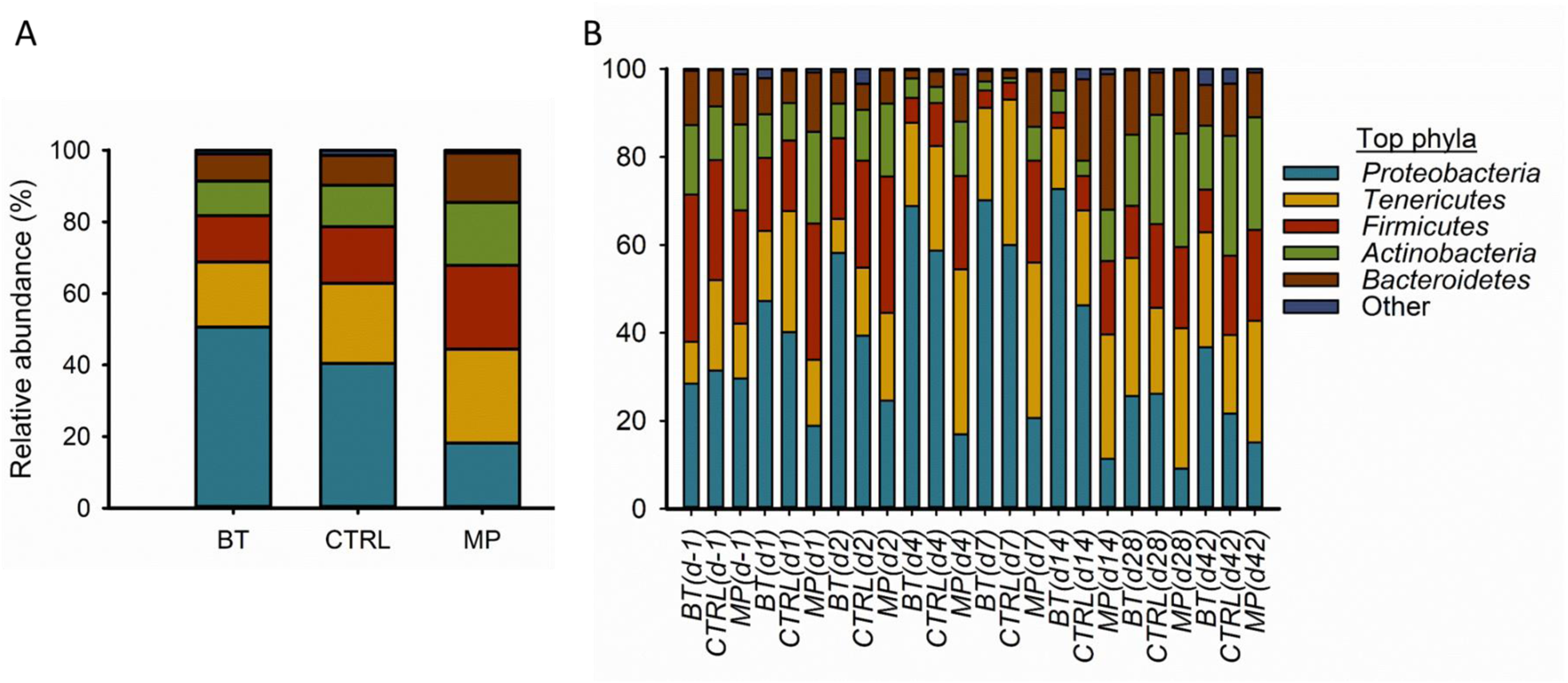
The five most relatively abundant phyla in the nasopharyngeal microbiota of calves. On day 1, cattle were treated with intranasal bacterial therapeutics (BT), intranasal PBS (CTRL), or subcutaneous tulathromycin (MP) (n = 20 per group). (A) overall comparison between treatment groups; (B) comparison by treatment groups and sampling day.

#### Changes in lactic acid-producing bacteria at the family level

The LAB taxonomically belong to the *Lactobacillales* order (28). Five different LAB families including *Aerococcaceae, Carnobacteriaceae, Enterococcacease, Lactobacillaceae* and *Streptococcaceae* were detected in the present study (Supplementary Fig. S2). Overall, no treatment effects on the abundance of these families were observed, although the relative abundance of *Lactobacillaceae* was >10% in four calves from the BT group.

#### Changes in microbial composition at genus level

Beta-binomial regression analysis allowed for identification of compositional differences of NP genera between treatment groups, across time. In total, we identified 28 genera within the BT and MP groups whose change in relative abundance from d-1 (baseline) was significantly different from changes that occurred in the CTRL group (*P* < 0.05). As shown in Fig. 6, 4 of the 10 most abundant genera (*Ruminococcaceae_UCG-005, Psychrobacter, Jeotgalicoccus* and *Planococcus*) were included among these taxa that differed between treatment groups. In general, most of the 28 taxa became less abundant in the BT group, following BT inoculation. Among which, *Ruminococcaceae_NK4A214_group*, *Paeniclostridium*, *Lachnospiraceae_NK4A136_group*, and *Cellvibrio* experienced a consistent decline during the entire post-BT inoculation period. For some taxa, the magnitude of change in relative abundance from the baseline varied among sampling times. For BT calves, the most significant reduction from baseline (d-1) was observed for *Ruminococcaceae_NK4A214_group* (d 2)*, Lachnospiraceae_NK4A136_group* (d 1), *Facklamia* (d 28), *Celllvibrio* (d 2) and *Acetitomaculum* (d 4). *Lactobacillus* was the only taxa in the BT group that experienced an increase in relative abundance that was >1, occurring on d 1. In contrast to BT group, most of the 28 genera became enriched in the MP calves, with several taxa having increases in relative abundances which were >2. The most immediate and consistent enrichment following tulathromcyin injection was observed for *Jeotgalibaca*.

**Fig. 6.**
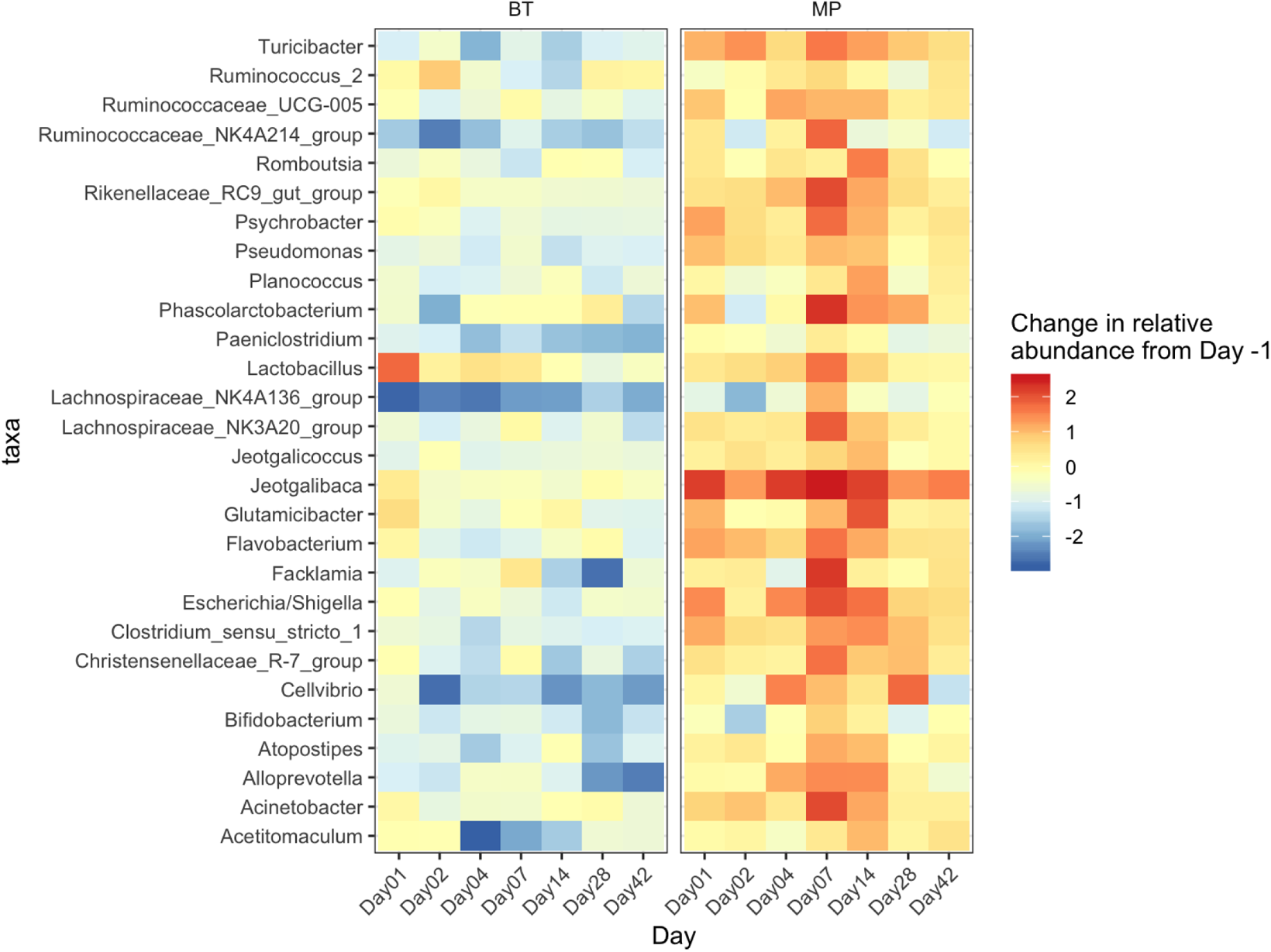
Heatmap comparing the bacterial microbiota in the nasopharynx of cattle. Taxa (n = 28) that showed a significant change (*P* < 0.05) from baseline (day −1) in BT and MP groups above and beyond any changes in the control (CTRL) group over the course of 42 days. On day 1, cattle were treated with intranasal bacterial therapeutics (BT), intranasal PBS (CTRL), or subcutaneous tulathromycin (MP) (n = 20 per group).

#### Changes in relative abundance of Lactobacillus spp

To identify whether the significant enrichment of genus *Lactobacillus* observed on d 1 (Fig. 2B) was due to the inoculated BT *Lactobacillus* strains, species-level taxonomic identification was performed on sequencing data from the *Lactobacillus* genus (Supplementary Fig. S3). Within *Lactobacillus*, 6 different taxonomic OTUs classified as *L. acetotlerans/fructivorans*, *L. acidophilud/amylovorus*, *L. amylovorus/buchneri*, *L. curvatus*/*graminis*, *L. fermentum/mucosae*, and *L. ruminis* were identified. Overall, the abundance of these OTUs varied however *L. acidophilus/amylovorus, L. amylovorus/buchneri, and L. curvatus*/*graminis* were generally only detected up to 48 h post-BT administration. This would suggest that the BT strains within the species *L. amylovorus*, *L. buchneri*, and *L. curvatus* accounted for the increase in these OTUs on d 1 and 2.

#### Changes in relative abundance of BRD-associated genera

The relative abundances of *Mannheimia, Pasteurella*, *Histophilus* and *Mycoplasma* which encompass BRD-associated pathogens, were not affected by treatment (*P* > 0.05; Fig. S4).

### Microbial interactions and dynamics of the NP microbiota

#### Interaction network structure among all observed OTUs

To evaluate overall dynamics of microbial communities, ecological modeling was used to analyse the interaction of all genera. As shown in network plots (Fig.7), distinct microbial interaction network structures were observed between CTRL, BT and MP groups. Compared to the CTRL group, the interaction network of microbiota from BT calves was more complex with a greater number of genera-genera interactions. In contrast, there was large decrease in genera-genera interactions among the microbial community of calves in the MP group, with only 22 genera identified in the network model. Even among these 22 genera, the interactions were connected by two separate hubs, indicating that the tulathromycin injection diminished the interaction network among the NP microbial community.

**Fig. 7.**
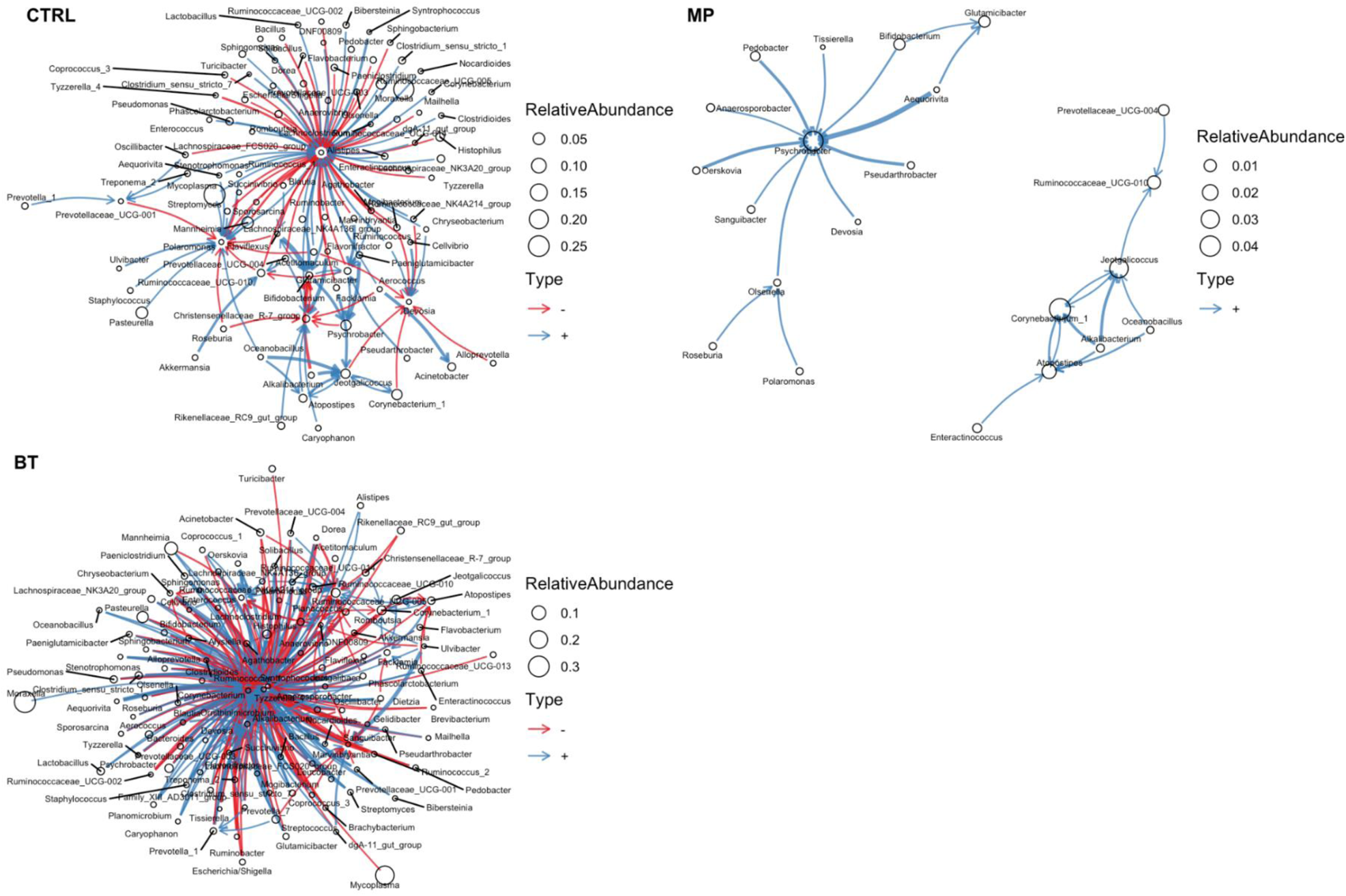
The ecological network of observed bacterial OTUs at the genus level in nasopharyngeal samples of calves. On day 1, cattle were treated with intranasal bacterial therapeutics (BT), intranasal PBS (CTRL), or subcutaneous tulathromycin (MP) (n = 20 per group).

#### Structure of causal relationships among 16 selected genera

Based on the observed distinct changes in species interaction network among NP microbiota in response to both BT and MP treatments, we further evaluated the relationship of 16 targeted genera using causal structure-based path modeling. This path model analysis provides greater detail on the interaction between OTUs, as it predicts causality and direction of the causality, and accounts for the time effect and unmeasured effects. Three path models (CTRL, BT, and MP) are described in Table 1 and are depicted as diagrams of causal relationships of the relative abundances in Fig. S5.

**Table 1.**
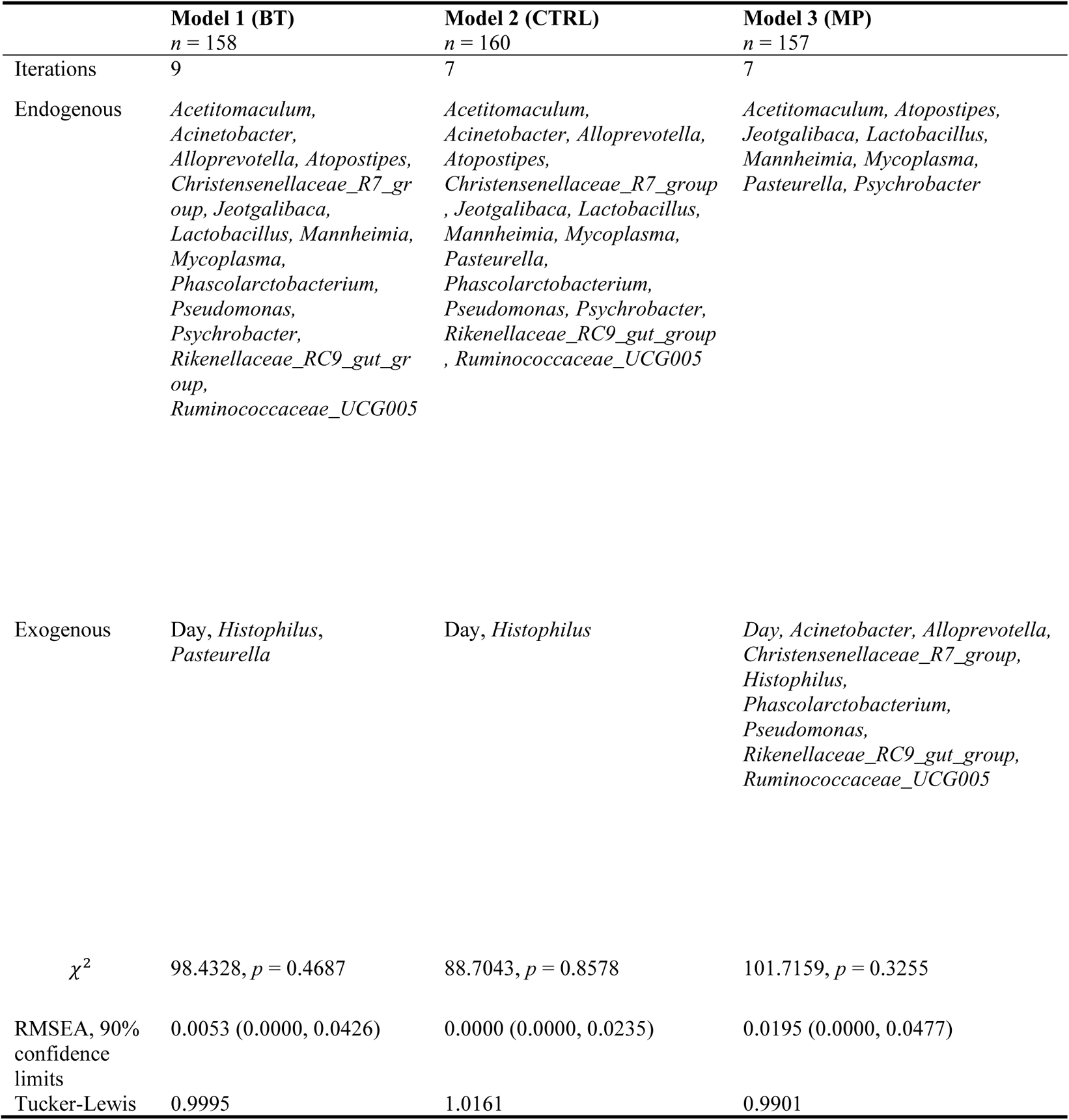
Manifest variables in the modified path models

|  | **Model 1 (BT)**<br>*n* = 158 | **Model 2 (CTRL)**<br>*n* = 160 | **Model 3 (MP)**<br>*n* = 157 |
| --- | --- | --- | --- |
| Iterations | 9 | 7 | 7 |
| Endogenous | *Acetitomaculum*,<br>*Acinetobacter*,<br>*Alloprevotella*, *Atopostipes*,<br>*Christensenellaceae_R7_group*, *Jeotgalibaca*,<br>*Lactobacillus*, *Mannheimia*,<br>*Mycoplasma*,<br>*Phascolarctobacterium*,<br>*Pseudomonas*,<br>*Psychrobacter*,<br>*Rikenellaceae_RC9_gut_group*,<br>*Ruminococcaceae_UCG005* | *Acetitomaculum*,<br>*Acinetobacter*, *Alloprevotella*,<br>*Atopostipes*,<br>*Christensenellaceae_R7_group*,<br>*Jeotgalibaca*, *Lactobacillus*,<br>*Mannheimia*, *Mycoplasma*,<br>*Pasteurella*,<br>*Phascolarctobacterium*,<br>*Pseudomonas*, *Psychrobacter*,<br>*Rikenellaceae_RC9_gut_group*,<br>*Ruminococcaceae_UCG005* | *Acetitomaculum*, *Atopostipes*,<br>*Jeotgalibaca*, *Lactobacillus*,<br>*Mannheimia*, *Mycoplasma*,<br>*Pasteurella*, *Psychrobacter* |
| Exogenous | *Day*, *Histophilus*,<br>*Pasteurella* | *Day*, *Histophilus* | *Day*, *Acinetobacter*, *Alloprevotella*,<br>*Christensenellaceae_R7_group*,<br>*Histophilus*,<br>*Phascolarctobacterium*,<br>*Pseudomonas*,<br>*Rikenellaceae_RC9_gut_group*,<br>*Ruminococcaceae_UCG005* |
| $\chi^2$ | 98.4328, *p* = 0.4687 | 88.7043, *p* = 0.8578 | 101.7159, *p* = 0.3255 |
| RMSEA, 90% confidence limits | 0.0053 (0.0000, 0.0426) | 0.0000 (0.0000, 0.0235) | 0.0195 (0.0000, 0.0477) |
| Tucker-Lewis | 0.9995 | 1.0161 | 0.9901 |

Details on model fit statistics and the number of iterations performed by the CALIS procedure are shown in Table 1. The *P*-values for the chi-square statistic for the modified path models were > 0.05, indicating good model fitting. The root mean square error of approximation (RMSEA) was < 0.05, and the 90% confidence limits of the RMSEA were also < 0.05. Therefore, the null hypothesis that the modified path models closely fit the data was retained. The values of the Tucker-Lewis index were also > 0.9, where the value of a true model would be expected to be 1.

The variables that constituted the subsets of exogenous (independent) and endogenous (dependent) variables varied in the three path models (Table 1). Day was initially hypothesized to be exogenous, and no path or covariance was added to change the position in the model of Day from being an independent explanatory variable. The process of model modification positioned the abundance of *Histophilus* as an exogenous variable in the modified path models, and it was a predictor variable with a statistically significant standardized total effect on *Alloprevotella*

(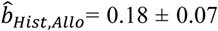, *P* = 0.005) in Model 1 (of the BT group) (Table S2), while it was not a predictor variable in Model 2 (CTRL) or Model 3 (MP) (Fig. S5).

The relative abundances of seven genera (*Acetitomaculum, Atopostipes, Jeotgalibaca, Lactobacillus, Mannheimia, Mycoplasma*, and *Psychrobacter*) were explained in the three modified path models. In Model 1 (BT group, Fig. S5A), *Pasteurella* was exogenous and therefore the sources of variance of *Pasteurella* abundance were outside of the model. In Model 2 (CTRL group, Fig. S5B), however, the variance of *Pasteurella* was explained by predictor variables: *Alloprevotella*, *Atopostipes, Christenseneelaceae_R7_group*, *Lactobacillus*, *Pseudomonas*, *Psychrobacter*, *Rikenellacease_RC9_gut_group*, *Ruminococcaceae_UCG005*, *Histophilus*, and Day.

#### Robust relationships

Path relationships that were statistically significant in the three modified models were considered to be robust. As such, the negative and statistically significant (all *P* <.0001) values of the standardized direct effect from Day to *Atopostipes* in the modified path models represent a robust causal relationship. Therefore, under the conditions of this experiment, the relative abundance of *Atopostipes* is expected to decrease monotonically over time. In contrast, the standardized direct effect from Day to *Psychrobacter* was positive in the modified path models, and therefore the relative abundance of *Psychrobacter* is expected to increase monotonically over time, under conditions similar to the experiment. Robust causal relationships were also identified between observed genera. For example, the standardized positive and statistically significant (*P* <.0001) total effects from *Psychrobacter* to *Jeotgalibaca* (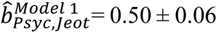; 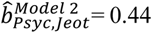 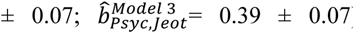) (Table S2); and from *Ruminococcaceae_UCG005* to *Acetitomaculum* (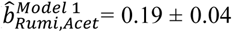; 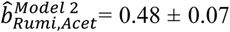; 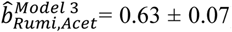) and *Jeotgalibaca* (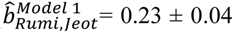; 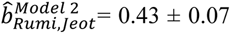; 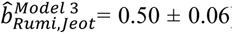). These positive path relationships can be interpreted as commensal biological relationships: where the bacteria that is a source of variance experiences no benefit or harm, but the bacteria that extracts variance can benefit by increasing monotonically in relative abundance.

#### Co-occurrence of path relationships for treatment groups

Path relationships among the observed genera were identified in Model 1 (BT group) and Model 2 (CTRL group). For the BT (Model 1) and CTRL (Model 2) groups, there were statistically significant (*P* ≤ .0001) and positive standardized total effects from *Christensenellaceae_R7_group* to *Rikenellaceae_RC9_gut_group* (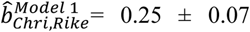; 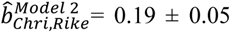) and from *Pseudomonas* to *Acetitomaculum* (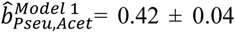; 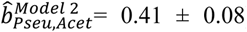), *Atopostipes* (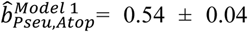; 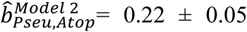), *Phascolarctobacterium* (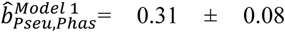; 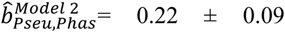), and *Rikenellaceae_RC9_gut_group* (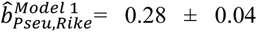; 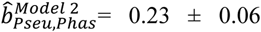). The standardized total effect from *Pseudomonas* to *Mannheimia* was negative (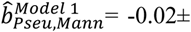 0.04, *P* = 0.69; 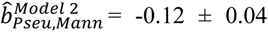, *P* = 0.002). Standardized total effects from *Psychrobacter* were positive and statistically significant (*P* = 0.022) to *Acetitomaculum* (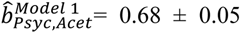; 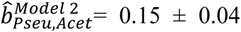), *Alloprevotella* (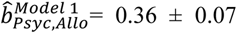; 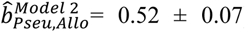), *Atopostipes* (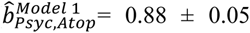; 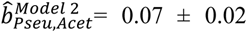), *Christensenellaceae_R7_group* (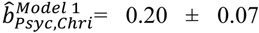; 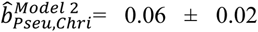), and *Rikenellaceae_RC9_gut_group* (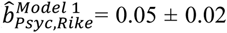; 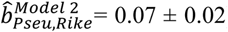). Furthermore, *Ruminococcaceae_UCG005* statistically significantly (*P* <.0001) and positively affected *Atopostipes* (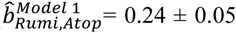; 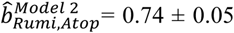), *Christensenellaceae_R7_group* (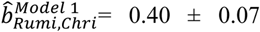; 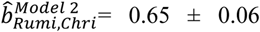), and *Rikenellaceae_RC9_gut_group* (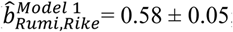; 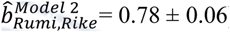).

In the MP group (Model 3, Fig. S5C), there were fewer path relationships that co-occurred in the BT (Model 1) and CTRL (Model 2) groups. The causal relationships between genera that were unidirectional for the CTRL and BT groups were frequently bi-directional in the MP group. The relative abundances of seven genera become exogenous variables in Model 3 (Table 1). One genus in the MP group (*Atopostipes*) exhibited standardized total effects that co-occurred in the BT group. The standardized total effects from *Atopostipes* were positive and statistically significant (*p* ≤ 0.02) to *Acetitomaculum* (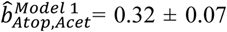; 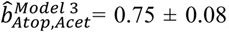) and *Jeotgalibaca* (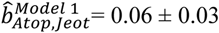; 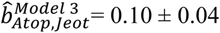).

The standardized total effects from the relative abundance of *Lactobacillus* to the relative abundances of other bacteria are shown in Table S3. In this case, the effects from *Lactobacillus* to *Acetitomaculum*, *Acinetobacter*, *Alloprevotella*, and *Jeotgalibaca* were positive in both BT and CTRL group. In Model 3 (MP), the relative abundance of *Lactobacillus* did not affect the relative abundances of the selected genera. Based on CTRL data (Model 2) *Lactobacillus* was modeled to increase monotonically the relative abundance of *Acetinomaculum*, which inhibits *Mannheimia* (Fig. S5). The effect of *Lactobacillus* on *Mannheimia* was therefore indirect in Model 2. In the BT group (Model 1), *Lactobacillus* had a direct positive effect and an indirect negative effect on *Mannheimia.* The indirect inhibition of *Mannheimia* was mediated by *Phascolarctobacterium*, which inhibited *Mannheimia*. The standardized total effects from *Lactobacillus* to *Mycoplasma* and *Pasteurella* were negative in the CTRL group, but these effects were not statistically significant in the BT group.

### Antimicrobial resistance determinants in the NP microbiota

The macrolide resistance gene *msr*(E) increased in the NP microbiome of MP calves during the first 28 days of the study (*P* < 0.01) (Fig. 8A). The abundance of *msr*(E) was greater in MP calves compared to CTRL and BT calves during the last two weeks of study (*P* < 0.05). The abundance of the tetracycline resistant gene, *tet*(H) was not affected by treatment or time (*P* > 0.05) (Fig. 8B).

**Fig. 8.**
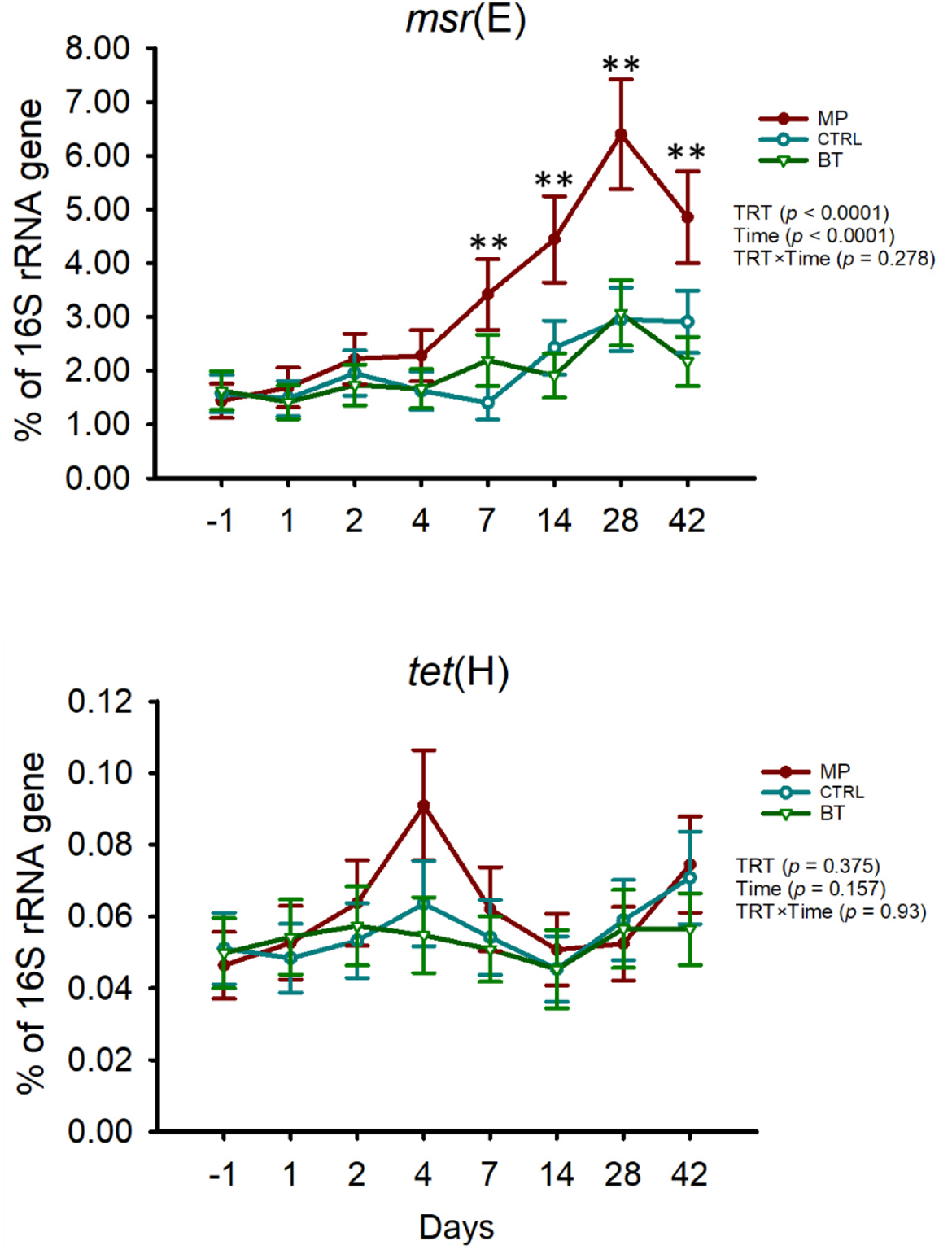
The proportion (%) of the resistance determinants msr(E) and tet(H) to 16S rRNA gene copies in nasopharyngeal samples obtained from cattle, quantified by qPCR. On day 1, cattle were treated with intranasal bacterial therapeutics (BT), intranasal PBS (CTRL), or subcutaneous tulathromycin (MP) (n = 20 per group). The results are presented as estimated mean ± standard error of the mean.

## Discussion

Studies have suggested that mutualistic and antagonistic interactions take place within the microbial community of the bovine respiratory tract (24, 25). These interactions may contribute positively or negatively to microbiota-mediated colonization resistance against respiratory pathogens (18, 29). Specifically, in the nasopharynx, LAB have been negatively associated with *Pasteurellaceae*, and certain LAB strains were capable of directly inhibiting BRD bacterial pathogens (25, 26). These data suggested that LAB have potential as BTs for mitigating BRD-associated pathogens. In the present study, we evaluated the effect of inoculating 6 previously characterized *Lactobacillus* strains (26) directly into the upper respiratory tract of cattle, on the microbiota, and compared those effects to a common metaphylactic antimicrobial, tulathromycin. Throughout the study, all calves remained healthy, with the exception of 5 that were treated for BRD. However, the rates of BRD were not affected by treatment, and weight gain was similar across all treatments, indicating that the BTs did not adversely affect inoculated calves.

### Colonization by BTs

A single application of BTs was selected, in order to fit within modern management systems of beef cattle, which employ a single processing event upon arrival at feedlots. Despite the BT strains originating from the nasopharynx of feedlot cattle and displaying strong *in vitro* adhesion to bovine turbinate cells (26), real-time PCR and 16S rRNA SV analyses indicated that colonization by BT strains was transient, lasting up to 48 h. Similar findings from gastrointestinal studies support the difficulty that exogenous strains have in colonizing established microbial communities (30, 31). The presence of similar indigenous species can affect colonization by exogenous bacteria (32), potentially by limiting resource availability (33, 34). For example, colonization by *Bifidobacterium longum* (AH1206) was impeded in the gastrointestinal tract when endogenous *B. longum* was already present (34). Indeed, *Lactobacillus* were detected prior to inoculation of BTs though these were mainly assigned to OTU *L. fermentum*/*mucosae*. Thus, inoculating the BTs earlier in the life of cattle may increase colonization potential if done prior to establishment of similar *Lactobacillus* spp.

### Longitudinal effects of BTs and tulathromycin on the respiratory microbiota

Changes in the composition of the NP microbiota were observed after BT administration. Despite BT treatment resulting in both increases and decreases of 28 taxa that significantly changed from baseline levels prior to inoculation (Fig. 6), the strongest BT effects on these taxa were inhibitory. The BT strains have previously been shown to produce lactate, hydrogen peroxide, or encode bacteriocins (26) which may have led to inhibition of resident bacteria. These direct impacts however would have only occurred within the 48 h the BTs colonizing the nasopharynx. It is interesting to note despite being transient, a single administration of the BTs had a long-term impact on structure and diversity of the NP microbiota. Both a reduction in richness and diversity was observed up to 42 days after BT inoculation. Information regarding the effect of probiotics or BTs on respiratory microbiota in animal models is limited, however similar effects have been reported for probiotics targeting the gastrointestinal microbiota. For example, Zhang and colleagues observed that the gut microbial community was impacted up to two weeks after administration of a *L. casei* probiotic strain (35). Other studies have reported prolonged effects on the gut microbiota for up to one (32) and five (36) months following cessation of a probiotic cocktail containing 11 strains fed to humans. The underlying mechanisms by which the BT *Lactobacillus* strains exerted long-term modulation of the microbiota are difficult explain but are likely related to their initial effects on community members.

Bacterial communities in most niches form complex ecological interaction webs, and such interactions are important in maintaining microbiome homeostasis and a symbiotic relationship between microbe and host (37). We therefore evaluated the microbiome wide community networks to gain insights into the changes in microbial community interactions in NP microbiota in response to BTs and antibiotic administration. The interaction network among all observed genera was predicted based on interaction networks from sequencing data. After observing the distinct interaction network structure among treatment groups, we decided to further investigate the causal networks among targeted taxa using structure equation modeling. Mainali et al. was the first to apply causal models to detect interaction networks in the human microbiome using conditional Granger causality (38). These authors argued that the causal models may provide more accurate prediction of interaction networks among microbial community relative to standard correlation/network analysis, as correlation is neither necessary nor sufficient to establish causation and environmental filtering can lead to correlation between non-interacting taxa.

The path model of the BT group revealed that a moderate degree of alterations had taken place in the causal relationship structure amongst observed genera, as seen by the changes in the magnitude of the direct and indirect effects from one genus to another. It is possible that the indirect effects of the BTs are what caused the prolonged effects on community diversity. Given the computational complexity of the path models, they were limited to 16 genera. However, it was interesting that ecological network analysis of all observed genera showed a more complex network for the BT calves compared to both CTRL and MP calves. Yang and colleagues suggested that probiotics (*Paracccus marcusii* DB11 and *Bacillus cereus* G19) promote intestinal microbiota homeostasis by enhancing species-species interactions and increasing the number of connecters and/or module hubs within the network (39). Thus, BT administration may have promoted NP microbiota homeostasis by strengthening and promoting species-species interactions. This in turn may have reduced colonization potential by new bacteria, which is supported by the decreased richness observed for BT calves. While we did not sample calves prior to arrival, several studies have shown that diversity of the NP microbiota increases within days after feedlot placement, and it has been suggested to be linked to susceptibility to BRD (19). It would therefore be interesting to measure the effect of BT inoculation in calves prior to transport in future studies, to evaluate whether they promote stabilization of the respiratory microbiota during transport.

Tulathromycin also altered the NP microbiota structure, diversity and composition, compared to CTRL calves. Measured by real-time PCR, the total bacteria per NP swab was reduced in MP calves on days 4 and 7. This was likely due to inhibition of members of the phylum *Proteobacteria*, which decreased in MP calves. Interestingly, despite this reduction, Shannon diversity and richness were increased in MP calves on d 7, compared to the other treatments. While Holman et al. (21) did not see an increase in diversity of the NP microbiota of calves administered tulathromycin, they did observe an increase in diversity in calves administered oxytetracycline. In addition, like our study, Holman and colleagues observed an increase in *Ruminococcaceae_UCG-005*, *Rikenellaceae_RC9_gut_group*, *Phascolarctobacterium*, *Facklamia*, *Jeotgalibaca*, and *Acinetobacter* following tulathromyin injection (21), supporting those injectable antimicrobials may lead to an increase in microbial richness and diversity of the respiratory microbiota.

Most often, the diversity of microbial community is claimed to be positively associated with the stability of microbiota (40) and health (41). Typically, however, studies showing positive associations between bacterial diversity and health related to the gastrointestinal tract (42, 43). However, it has also been argued that diversity in host-associated microbial communities may not always be associated with microbial community stability and health. For example, higher bacterial diversity and richness were observed in upper respiratory tract of children with invasive pneumococcal disease compared to healthy children (44). It was proposed by the authors that the higher diversity and richness of the NP microbiota was associated with impaired immune response. It was interesting in our study that antimicrobial treatment increased diversity of the respiratory tract, whereas antimicrobial administration can reduce diversity in the gastrointestinal tract. Perhaps this is reflection of increased exposure to exogenous bacteria that the respiratory tract faces compared to the digestive tract.

Functional properties of the gut microbiota including colonization resistance at the community level are believed to be maintained by the active interactions among genetically distinct and diverse microbial species, which allows the microbial community to perform complex metabolic activities. A single mono-species population or multi-species but with no interconnectivity could not provide such collective functions of microbiota (45). In addition, the functional activities and stability of a microbiota is influenced by the positive and negative feedback loops generated as a result of the cooperation (46) and competition (47) among the different microbial species (48). Also, evolution of species-species interactions has been reported to determine microbial community productivity in new environments (49). Although proportionally equal positive and negative interactions between the bacterial species within NP microbiota were observed in both CTRL and BT groups, such cooperative and competitive interactions were more intensive in BT group compared to CTRL group. In contrast, there was only positive interactions observed between these fewer species that remained in network model as interconnected species in the MP group.

The stability of a mammalian microbiota depends on how the species interacts with one another (50). Weak and competitive interactions are stabilizing, and they limit positive feedback loops and the possibility that, if one species decreases, it will result in the decrease of others. Cooperative interactions determine the productivity of microbiome which is the efficiency of converting resources into energy (50). In the MP group, there was only a cooperative interaction observed in NP microbiota and the competitive interaction was missing, suggesting that NP microbiota in MP cattle would most likely experienced dysbiosis. Whereas, in BT calves, both cooperative and competitive interactions were present with higher magnitude. This indicated that the NP microbiota in BT calves was most likely stable with normal community functions. It has been argued that the asymmetry of an unhealthy microbiome can relate to non-neutral states created by strong stressors, reducing host ability to contain certain bacteria and resulting in overgrowth of abundance and multiplication of species (51). We observed that the abundance of most of the observed significant taxa (n = 28) were enriched in calves that received antibiotic. The overgrowth of these taxa therefore might be due to the non-neutral state of microbiota induced by antibiotic.

While the increase in diversity following tulathromcyin was unexpected, we hypothesize that it was related to the effects of the antimicrobial on the community network. For MP calves, 7 genera were determined to be exogenous to the developed path model, which had fewer interactions than those developed for the BT and CTRL groups of calves. In addition, the ecological network was also far less complex, and together these data indicate a substantial perturbation of the ecological network in MP calves. Likewise, Yang et al. (39) reported that the use of the antimicrobial florfenicol resulted in deterioration of the ecological network among intestinal microbiota of sea cucumbers, leading to the homeostatic collapse of microbiota. Thus, the deterioration of the species interactions by tulathromycin may have made the NP microbiota in MP calves more permissive to exogenous bacteria colonization, increasing diversity.

While the impact this may have on cattle health was not possible to evaluate in the present study, it is an area that warrants further investigation. Despite the use of metaphylactic antimicrobials, the overall rate of BRD in feedlots has not declined over the last 40 years (4). In fact, some feedlots are observing a change in BRD incidence, with cases occurring later in the feeding period (communication: Dr. Eugene Janzen, University of Calgary). While unsupported, it is tempting to speculate that metaphylaxis treatment may prevent short-term BRD incidence through direct inhibition of pathogens, but the causal deterioration of keystone taxa (52) may increase risk of longer-term respiratory complications. In contrast, we observed that BTs promoted microbiome homeostasis by stabilizing the NP bacterial community structure and genera interactions.

### Longitudinal effects of BTs and tulathromycin on BRD-associated pathogens

The MP treatment reduced the prevalence of culturable *M. haemoltyica*. In support of this, tulathromycin injection has previously been shown to reduce NP colonization of *M. haemolytica* in feedlot cattle (10). In contrast, despite showing strong inhibitory properties against *M. haemotytica in vitro*, the BTs did not reduce prevalence of *M. haemolytica.* However, the BT strains were tested for competitive exclusion *in vitro* (26). The fact that 24% of calves in the BT group were *M. haemoltyica*-positive prior to administration suggests that the BT strains may not be effective for displacement of *M. haemolytica*.

Despite MP calves being the only group to have several time points where no *M. haemolytica* or *P. multocida* could be cultivated from NP swabs, sequence analysis of *Mannheimia*, *Pasteurella*, *Histophilus,* and *Mycoplasma* revealed no differences in relative abundance of these BRD-associated genera. For *Mannheimia*, *Pasteurella*, and *Histophilus*, this was likely a result of inter-animal variation and overall limited relative abundance of these genera. While a previous study did show that tulathromycin injection reduced NP *Pasteurella*, the calves in that study were ranch-derived and had higher levels of *Pasteurella* at feedlot entry (21). The limited number of BRD cases and reduced relative abundance of BRD-associated genera in our study was likely attributed to the calves being at lower risk for BRD, having come from a single source and being placed into individual stalls instead of being co-mingled.

It was interesting that *Histophilus* was exogenous to microbial path models for all three treatments. *H. somni* has previously been shown to have a 32.5 times greater chance of being isolated from feedlot calves after 40 days on feed (14). This suggests that genera that were not included in the models, or external factors associated with feedlots, may be related to its colonization of cattle. *Lactobacillus* had a direct positive effect on *Mannheimia*, for BT calves, though indirect negative effects were also observed. This finding is difficult to explain but may be related to limiting model analysis to 16 genera. Although the relative abundance of *Mannheimia* did not increase as a result of BT treatment, future studies with *Lactobacillus*-based BTs should be tested for their effects on *Mannheimia*. In contrast, for CTRL calves, *Lactobacillus* had overall negative effects on *Mannheimia* and *Pasteurella* though they were not direct. Thus, this supports previous data showing the importance of indigenous *Lactobacillus* in bovine respiratory health (22, 24).

### Longitudinal effects of BTs and tulathromycin on antimicrobial resistant determinants

The macrolide resistance gene *msr*(E) encodes a multidrug efflux pump conferring resistance to macrolides, including tulathromycin (53). This gene has been detected in BRD pathogens and has been detected within integrative conjugate elements (15, 16). Given the increase in macrolide resistance in feedlot BRD pathogens over the last 10 years (11, 14), evaluation of resistance genes in respiratory bacteria is important, to maintain proper selection of antimicrobials. Despite altering the microbiota, BT-treated calves did not select for bacteria that encoded *msr*(E) or *tet*(H). In contrast, *msr*(E) increased in MP calves showing that tulathromycin administration selected for bacteria carrying this resistance gene. Similarly, tulathromycin use has previously been associated with increased antimicrobial resistance determinants in NP microbiomes of feedlot cattle (21, 54).

## Conclusion

A single dose of intranasal BTs, which were developed from bovine respiratory commensal *Lactobacillus* spp., induced longitudinal modulation of the NP microbiota in auction market beef calves, with no adverse effects on animal health and growth performance. While no differences in the relative abundances of BRD-associated genera were observed between treatments, the ecological networks of NP bacteria from BT-treated calves became more integrated. It was proposed that this resulted in a more stable microbiome with increased resilience against exogenous microorganisms. In contrast, disruption of the microbiome after tulathromycin treatment reduced resilience, leading to increased diversity in MP calves. Overall, this study showed that the bovine respiratory microbiota can be altered by administration of BTs and may therefore provide new opportunities to enhance microbiome-mediated respiratory resistance against BRD pathogens. We also showed the usefulness of employing network and structure equation modelling to evaluate the impact of BTs on host microbiota. Future studies should consider the optimal administration of BTs, as improved resilience against BRD pathogens may result from administration prior to calves being shipped to feedlots.

## Materials and Methods

### Animals and experimental design

Animals used in this study were cared for in agreement with the Canadian Council for Animal Care guidelines (55). All the procedures and protocols with respect to animal handling and sampling were reviewed and approved by the Animal Care Committee at the Lethbridge Research and Development Centre, Agriculture and Agri-Food Canada (Lethbridge, Alberta, Canada).

Sixty crossbred beef heifers approximately 6 months-old (initial BW = 266 ± 13 kg) and originating from a single cow-calf ranch, were purchased from a local auction market and transported to the Lethbridge Research and Development Centre feedlot (<10 km distance). Upon arrival, the calves were weighed, and NP swabs were collected (d −1). Calves were then blocked by weight and randomly assigned to three treatment groups (*n* = 20 per treatment): i) BT group received an intranasal cocktail of six *Lactobacillus* strains suspended in phosphate buffered saline (PBS) in equal concentrations (3 × 10^9^ CFU per nostril), ii) metaphylaxis (MP) group received a subcutaneous injection of tulathromycin (2.5 mg/kg BW), and iii) control (CTRL) group received intranasal PBS without bacteria. The calves were housed individually in pens throughout the study and were fed once daily a diet containing 75% barley silage, 22.5% dry roll barley, and 2.5% standard feedlot supplement. Calves had free access to drinking water.

### Preparation of BT inoculum

The BT cocktail was a mixture of six *Lactobacillus* strains (1×10^9^ CFU mL^−1^): *L. amylovorus* (isolate 72B), *L. buchneri* (63A and 86D), *L. curvatus* (103C) and *L. paracasei* (3E and 57A). These isolates were inoculated on *Lactobacillus* De Man, Rogosa and Sharpe (MRS) agar (Dalynn Biologicals, Calgary, AB, Canada), and incubated for 48 h at 37°C in 10% CO_2_. One day prior to nasal inoculation, a single colony of each strain was inoculated into 5 mL MRS broth and incubated at 37°C with agitation at 200 rpm. After 18 h of incubation, each bacterial culture was centrifuged at 7,600 × *g* for 10 min, the supernatant discarded and the pellet re-suspended with pre-warmed (37°C) PBS to achieve a target concentration of 1 ×10^9^ CFU per mL^−1^ using pre-established OD_600_ values. The BT cocktail was prepared by mixing the six *Lactobacillus* isolates at equal ratios in PBS. One hour prior to inoculation, 3 mL of the BT cocktail was loaded in a sterile 10 mL syringe and the syringe tip was covered with sterile needle to prevent any leakage. For the control group, 3 mL PBS was loaded similarly into a syringe.

### Administration of BT cocktail and tulathromycin

On day 1, calves were restrained in a squeeze chute and administered treatments. Sterile laryngo-tracheal mucosal atomization devices (LMA® MADgic® Laryngo-Tracheal Mucosal Atomization Device without syringe, Cat# MAD 700, Teleflex, Morrisville, NC) were fitted to loaded syringes, and the atomization device was inserted into each nostril of calves (approximately 15 cm) and sprayed until the syringe was empty. One atomization device was used for the two nostrils of each calf. A total of 6 mL of BT inoculum (3 mL per nasal cavity) was administrated to calves in the BT treatment group. For control group animals, PBS was sprayed into the nostrils similar to the BT inoculum (3 ml per nostril). The metaphylaxis group received a single subutaneous injection of long-acting tulathromycin (2.5 mg/kg body weight).

### Nasopharyngeal swab sampling and processing

In addition to sampling at feedlot arrival (d −1), NP samples were collected from the right nostril of each calf in the study on days 1, 2, 4, 7, 14, 28, and 42. The NP sampling procedures were described previously (19). Prior to sampling, the right nostril was wiped clean with 70% ethanol. Extended guarded swabs (27 cm) with a rayon bud (MW 124, Medical Wire & Equipment, Corsham, England) were used for sampling. Swabs were taken while the animals were restrained in a squeeze chute. Swab tips were then cut and placed in a sterile 1.5 mL tube on ice. Samples were transported to the lab and processed within one hour of collection. At the lab, the swab tip was transferred into a cryovial containing 1 mL brain heart infusion (BHI) with 20% glycerol and vortexed.

#### Isolation and detection of bovine respiratory pathogens

Aliquots of swab suspension were plated for isolation and detection of BRD-associated pathogens including *M. haemolytica, P. multocida*, and *H. somni.* Culturing, isolation, and PCR identification of the pathogenic isolates were described previously (19, 22). The remaining swab suspensions in BHI glycerol stock were stored at −80 °C for DNA extraction.

### Genomic DNA extraction, 16S rRNA gene sequencing and analysis

The genomic DNA was extracted from the swab suspension according to the methods of Holman et al. (19, 21). From the extracted DNA, the V4 region of the 16S rRNA gene was amplified using primers 515-F (5′-GTGYCAGCMGCCGCGGTAA-′3) and 806-R (5′-GGACTACNVGGGTWTCTAAT-′3) (21). The amplicon was sequenced on a MiSeq instrument (Illumina, San Diego, CA, USA) with the MiSeq Reagent Kit v2.

After quality check with FastQC 0.11.5 and MultiQC 1.0 (56) primers and low quality sequences were trimmed off the raw sequence reads using cutadapt 1.14 (57). The trimmed reads were used to construct amplicon sequence variants (ASVs) using dada2 1.10.0 (58) in R 3.5.1 (59). Unless otherwise stated, all dada2 functions were used with default parameters. Reads were first filtered with dada2::filterAndTrim with a max-expected error of 1. Error rates were learned for the forward and reverse reads separately and these error rates were used to infer exact sequences (error correct) for each sample from dereplicated, trimmed reads using pooled=TRUE for the dada2::dada. Following this, the forward and reverse reads were merged using dada2::mergePairs. Chimeras were removed with dada2::removeBimeraDenovo and taxonomy was assigned using the naïve Bayesian classifier (60) as implemented in dada2::assignTaxonomy trained with the Silva training set version 132 (https://doi.org/10.5281/zenodo.1172782). Species level assignment was done with dada2::addSpecies that uses exact matching to assign species where possible. ASVs were aligned with ssu-align 0.1.1 (61). Quantification of bacterial, *Lactobacillus* spp., and antibiotic resistance determinants using quantitative PCR.

Real-time PCR was performed to quantify copies of 16Sr RNA genes in DNA from nasal swabs that were specific to all bacteria, or *Lactobacillus*, to estimate the abundances of total bacteria and *Lactobacillus* genera, respectively. Total 16S rRNA gene copies were amplified using primers 515F and 806R described above. *Lactobacillus*-specific 16S rRNA gene copies were amplified using a genus-specific primer described previously (62). In addition to 16s rRNA genes, the macrolide resistance gene *msr*(E) and tetracycline resistance gene *tet*(H) were quantified from the DNA extracted from nasal swabs. The primers for *msr*(E) and *tet*(H) were reported previously by Klima et al. (15) and Zhu et al. (63), respectively.

To generate standards for PCR, amplicons of each target gene were cloned into competent *E.coli* cells using the TOPO Cloning Reaction Kit (Invitrogen) according to the instructions of the manufacturer. Plasmids containing amplicon inserts were purified by the QIAprep spin miniprep kit (Qiagen, Hilden, Germany) and then serially diluted. Each real-time PCR mixture (25 μL) contained 1X iQ SYBR Green Supermix (Bio-Rad Laboratories Inc.), 0.4 μM of each primer, 0.1 μg/μL BSA (New England Biolabs,180 Pickering, ON, Canada), and 25 ng of DNA. For each PCR reaction, the amount of DNA extracted from the NP swabs was normalized to 10 ng/µL. The quantification of target genes was performed on a CFX96 Touch Real-Time PCR Detection system (Bio-Rad Laboratories Inc.) with the following conditions: an initial denaturation at 95°C for 3 min, followed by 40 cycles at 95°C for 25 sec, 50°C for 30 sec, and then 72°C for 45 s. For quantification of total and *Lactobacillus*-specific 16S rRNA gene copies and the resistant gene copies, standards were prepared for each gene using the respective p-Drive plasmid containing inserted amplicons and concentrations of 10^6^, 10^5^, 10^4^, 10^3^, and 10^2^ copies per reaction (in duplicate). Melt curve analysis was performed on all PCR reactions to ensure specific amplification. The temperature range was 60°C to 95°C and fluorescence was measured at 0.5°C intervals.

### Statistical analysis

Statistical analysis of sequence data was done with R 3.6.0 with phyloseq 1.28.0 (56), and vegan 2.5.5 (57). Plots were created with ggplot2 3.1.1. Sequences matching mitochondria or chloroplast were removed along with any sequences that were not assigned to Bacteria. A filtered copy of the ASV sequence table was created that retained ASVs present (count ≥ 2) in at least 1% of the samples. This served to reduce noise for downstream analysis. The full version of the sequence table was used for alpha diversity, which was assessed with the Shannon diversity index using 0.3.1 (64). Richness was estimated with a Poisson model using breakaway 4.6.8 (65). To calculate beta diversity, the ASV counts (filtered table) were normalized by with a variance stabilizing transform (using DESeq2 1.24.0) (66) with size factors calculated using GMPR (67) then sample-sample distances were determined with the Bray-Curtis metric and visualized with detrended correspondence analysis (DCA). Permutational multivariate analysis of variance (PERMANOVA) with 10,000 permutations was used to determine the effect of treatment, and sampling time on the microbial community structure. For statistical testing, the model included an interaction between day and treatment to detect differences between treatment groups over time. The betta function from breakaway 4.6.8 (68), which models both observed and unobserved diversity, was used to test alpha diversity. Corncob 0.1.0 (69) was used to test for differentially abundant taxa by fitting a model with and without the interaction term. This identified taxa that showed a change from baseline in the BT and MP groups differed significantly from the CTRL group. Corncob uses a beta-binomial-based model that controls for correlated observations (taxa) and overdispersion.

Generalized linear models (SAS PROC GLIMMIX) were built on the prevalence data of *M. haemolytica*, *P. multocida* and *H. somni* determined by culturing. The Bernoulli/binary distributions were selected for the models. Separate *F*-tests of the treatment effect were produced, using the SLICE option in the LSMEANS statement, for the respective days of observations.” The relative abundance of phyla and qPCR results data was analyzed using the GLIMMIX procedure in SAS (SAS 9.4, SAS Institute Inc., Cary, NC). The individual calf, treatment, and time were included in the CLASS statement. The models were “generalized” due to the specification of response distributions that were not Gaussian normal. Models were “mixed” due to the inclusion of fixed effects (treatment-nested-in-time and time) and random effects (individual). Variance heterogeneity was modeled using a “RANDOM _RESIDUAL_ / GROUP = Treatment*Time” statement. Response distributions and structures of the variance-covariance matrix were selected for each genus based on the model fit statistics, i.e., the Bayesian information criterion (BIC). Preliminary models that specified the beta-binomial distribution did not converge. Therefore, alternative distributions were tested: Gamma, inverse Gaussian, lognormal, shifted *t*, Gaussian normal, exponential, and geometric.

Ecological network modeling was performed to evaluate the directed microbial interactions among the nasopharyngeal microbial communities using BEEM-static in R. Briefly, based on the abundance profile of all observed genera derived from 8 sampling time points, generalised Lotka-Volterra models (gLVMs) coupling biomass estimation and model inference in an expectation maximization-like algorithm (BEEM) was used to construct three (CTRL, BT, MP groups) ecological network models as described by Li et al. (70). The interaction network inferred by BEEM-statistic was visualized by the plots generated using graph package of R.

Path analysis was performed to model the interrelationships of selected bacterial species within the NP microbial communities. Path analysis is a member of the structural equation modeling tools that enables the identification of causal relationships between measured variables (71). The construction of an initial path model was based on theories regarding the causal relationships. These path diagrams were used to illustrate the strength and direction of the causal relationships between variables. To identify biological interactions within the NP microbiota, and the changes in biological interactions in response to the BT and MP treatment, 16 genera were selected: 12 that exhibited the greatest change in relative abundance in BT and MP groups relative to CTRL group over the course of study, and four BRD-associated genera (*Mannheimia*, *Pasteurella*, *Histophilus*, and *Mycoplasma*). The relative abundance data for these 16 genera were sorted into three subsets based on the treatment groups (BT, CTRL and MP). The methods of path modeling followed those of Schwinghamer et al. (71). The CORR procedure in SAS (SAS 9.4, SAS Institute Inc., Cary, NC, USA) was used to calculate the matrices of Spearman rank-based correlation coefficients that were used as input for path modeling with SAS PROC CALIS. The initial hypothetical model was:

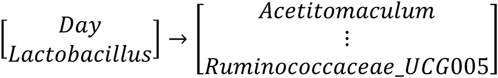

Three modified path models were developed based on the addition and subtraction of paths and covariance terms, based on Wald statistics and Lagrange multiplier statistics that were calculated using the MODIFICATION option in the PROC CALIS statement. Modifications were selected based on lower (better) values of Schwarz’s BIC and a value equal to zero for the stability criterion of reciprocal causation.

## Acknowledgement

The authors would like to thank Dr. Alexander’s lab team members including Dr. Long Jin, Pamela Caffyn, and Leandra Schneider, as well as the feedlot crew at the Lethbridge Research and Development Centre for their technical support during the animal trial.

## Funding

The work presented in this study was supported by Results Driven Agricultural Research (RDAR; Project #2016R032R). SA was the recipient of the Canadian Natural Science and Engineering Research Council (NSERC) Doctoral Scholarship.

## Consent for publication

Not applicable.

## Competing interests

The authors declare that they have no competing interest

---

**Supplementary Table S1.**
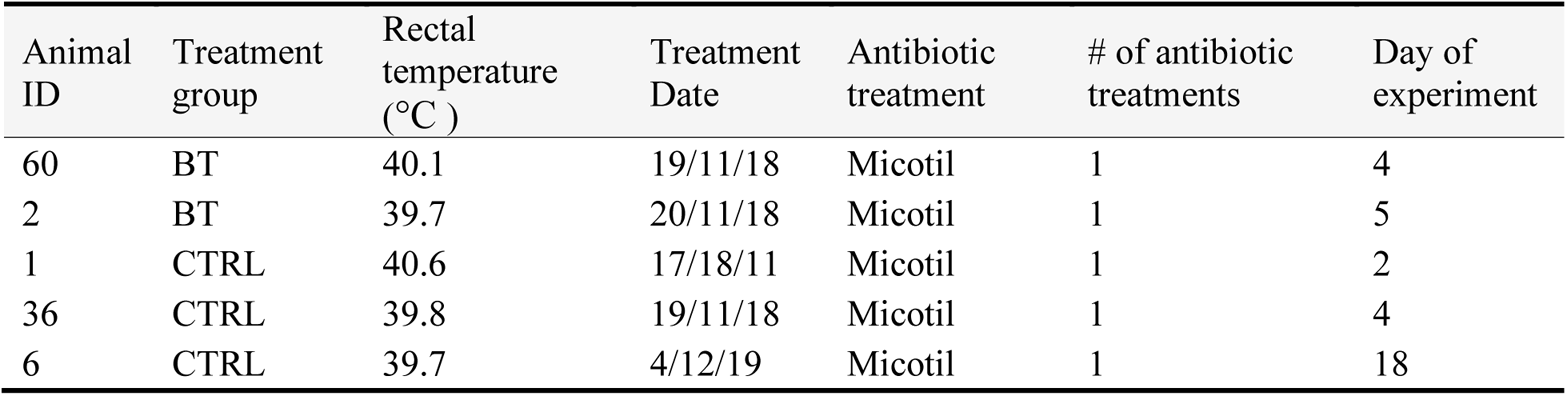
Animal health records

| Animal ID | Treatment group | Rectal temperature (°C ) | Treatment Date | Antibiotic treatment | # of antibiotic treatments | Day of experiment |
| --- | --- | --- | --- | --- | --- | --- |
| 60 | BT | 40.1 | 19/11/18 | Micotil | 1 | 4 |
| 2 | BT | 39.7 | 20/11/18 | Micotil | 1 | 5 |
| 1 | CTRL | 40.6 | 17/18/11 | Micotil | 1 | 2 |
| 36 | CTRL | 39.8 | 19/11/18 | Micotil | 1 | 4 |
| 6 | CTRL | 39.7 | 4/12/19 | Micotil | 1 | 18 |

**Supplementary table S2.** Standardized total effect (mean ± SE) from predictor variables (listed on the columns) on the dependent variables (listed on the rows) (Model 1, BT group). **Supplementary table S2.** Continue (Model 2, CTRL group). **Supplementary table S2.** Continue (Model 3, MP group).

|  | Ace | Aci | Ato | Chr | Lac | Man | Pha | Pse | Psy | Rum | Day | His | Pas |
| --- | --- | --- | --- | --- | --- | --- | --- | --- | --- | --- | --- | --- | --- |
| **Ace** | 0 | 0.10 ±<br>0.025<br>$p < .0001$ | 0.32 ±<br>0.07<br>$p < .0001$ | 0 | 0.24 ±<br>0.009<br>$p = 0.0073$ | 0 | 0 | 0.42 ±<br>0.044<br>$p < .0001$ | 0.68 ±<br>0.053<br>$p < .0001$ | 0.19 ±<br>0.042<br>$p < .0001$ | -0.05 ± 0.051<br>$p = 0.3016$ | 0 | -0.08 ± 0.026<br>$p = 0.0034$ |
| **Aci** | 0 | 0 | 0 | 0 | 0.24 ±<br>0.062<br>$p = 0.0001$ | 0 | 0 | 0.08 ±<br>0.026<br>$p = 0.0038$ | 0 | 0 | -0.02 ± 0.007<br>$p = 0.0405$ | 0 | 0 |
| **All** | 0.20 ±<br>0.081<br>$p =$<br>0.0116 | 0.11 ±<br>0.027<br>$p = 0.0001$ | 0.07 ±<br>0.029<br>$p = 0.0284$ | 0 | 0.03 ±<br>0.009<br>$p = 0.0076$ | 0 | 0.18 ±<br>0.072<br>$p = 0.0135$ | 0.28 ±<br>0.045<br>$p < .0001$ | 0.36 ±<br>0.072<br>$p < .0001$ | 0.10 ±<br>0.029<br>$p = 0.0005$ | -0.25 ± 0.068<br>$p = 0.0001$ | 0.18 ±<br>0.065<br>$p = 0.0048$ | -0.04 ± 0.016<br>$p = 0.0099$ |
| **Ato** | 0 | 0.13 ±<br>0.031<br>$p < .0001$ | 0 | 0 | 0.03 ±<br>0.011<br>$p = 0.0064$ | 0 | 0 | 0.54 ±<br>0.042<br>$p < .0001$ | 0.88 ±<br>0.05<br>$p < .0001$ | 0.24 ±<br>0.052<br>$p < .0001$ | -0.41 ± 0.053<br>$p < .0001$ | 0 | -0.10 ± 0.034<br>$p = 0.0034$ |
| **Chr** | 0 | 0.28 ±<br>0.044<br>$p < .0001$ | 0.22 ±<br>0.075<br>$p = 0.0029$ | 0 | 0.10 ±<br>0.025<br>$p = 0.0002$ | 0.14 ±<br>0.053<br>$p = 0.0087$ | 0.24 ±<br>0.064<br>$p = 0.0002$ | 0.34 ±<br>0.043<br>$p < .0001$ | 0.20 ±<br>0.067<br>$p = 0.0033$ | 0.40 ±<br>0.067<br>$p < .0001$ | -0.34 ± 0.055<br>$p < .0001$ | 0 | -0.02 ± 0.011<br>$p = 0.0034$ |
| **Jeo** | 0 | 0.15 ±<br>0.029<br>$p < .0001$ | 0.06 ±<br>0.025<br>$p = 0.0168$ | 0.27 ±<br>0.066<br>$p < .0001$ | 0.01 ±<br>0.02<br>$p = 0.5419$ | -0.12 ±<br>0.063<br>$p = 0.0632$ | 0.10 ± 0.03<br>$p = 0.0005$ | 0.37 ±<br>0.043<br>$p < .0001$ | 0.50 ±<br>0.058<br>$p < .0001$ | 0.23 ±<br>0.041<br>$p < .0001$ | -0.01 ± 0.045<br>$p = 0.904$ | 0 | -0.06 ± 0.020<br>$p = 0.0047$ |
| **Lac** | 0 | 0 | 0 | 0 | 0 | 0 | 0 | 0.32 ±<br>0.072<br>$p < .0001$ | 0 | 0 | -0.06 ± 0.026<br>$p = 0.0137$ | 0 | 0 |
| **Man** | 0 | -0.08 ±<br>0.03<br>$p = 0.0118$ | 0 | 0 | 0.19 ±<br>0.076<br>$p = 0.0134$ | 0 | -0.26 ±<br>0.075<br>$p = 0.0004$ | -0.02 ±<br>0.039<br>$p = 0.6922$ | 0 | 0 | 0.003 ±<br>0.008<br>$p = 0.6948$ | 0 | 0 |
| **Myc** | 0 | 0 | 0 | 0 | 0 | 0 | 0 | 0 | 0 | 0 | 0.33 ± 0.071<br>$p < .0001$ | 0 | 0 |
| **Pha** | 0 | 0.29 ±<br>0.078<br>$p = 0.0002$ | 0 | 0 | 0.07 ±<br>0.026<br>$p = 0.0077$ | 0 | 0 | 0.31 ±<br>0.075<br>$p < .0001$ | 0 | 0 | -0.06 ± 0.025<br>$p = 0.0164$ | 0 | 0 |
| **Pse** | 0 | 0 | 0 | 0 | 0 | 0 | 0 | 0 | 0 | 0 | -0.20 ± 0.066<br>$p = 0.0026$ | 0 | 0 |
| **Psy** | 0 | 0.15 ±<br>0.035<br>$p < .0001$ | 0 | 0 | 0.04 ±<br>0.013<br>$p = 0.0063$ | 0 | 0 | 0.62 ±<br>0.043<br>$p < .0001$ | 0 | 0.28 ±<br>0.056<br>$p < .0001$ | 0.19 ± 0.057<br>$p = 0.0008$ | 0 | -0.11 ± 0.038<br>$p = 0.0029$ |
| **Rik** | 0 | 0.32 ±<br>0.044<br>$p < .0001$ | 0.06 ±<br>0.024<br>$p = 0.021$ | 0.25 ±<br>0.069<br>$p = 0.0002$ | 0.08 ±<br>0.023<br>$p = 0.0002$ | 0.04 ±<br>0.016<br>$p = 0.033$ | 0.06 ±<br>0.023<br>$p = 0.0095$ | 0.28 ±<br>0.039<br>$p < .0001$ | 0.05 ±<br>0.022<br>$p = 0.0216$ | 0.58 ±<br>0.054<br>$p < .0001$ | -0.12 ± 0.032<br>$p = 0.0001$ | 0 | -0.01 ± 0.003<br>$p = 0.0684$ |
| **Rum** | 0 | 0.53 ±<br>0.054<br>*p* < .0001 | 0 | 0 | 0.13 ±<br>0.035<br>*p* = 0.0004 | 0 | 0 | 0.40 ±<br>0.056<br>*p* < .0001 | 0 | 0 | -0.08 ± 0.029<br>*p* = 0.0062 | 0 | 0 |

|  | **Ace** | **All** | **Ato** | **Chr** | **Jeo** | **Lac** | **Pse** | **Psy** | **Rik** | **Rum** | **Day** | **His** |
| --- | --- | --- | --- | --- | --- | --- | --- | --- | --- | --- | --- | --- |
| **Ace** | 0 | 0.08 ± 0.026<br>*p* = 0.0020 | 0 | 0.08 ± 0.032<br>*p* = 0.014 | 0.28 ± 0.073<br>*p* = 0.0001 | 0.30 ± 0.100<br>*p* = 0.0021 | 0.41 ± 0.070<br>*p* < .0001 | 0.16 ± 0.037<br>*p* < .0001 | 0 | 0.48 ± 0.055<br>*p* < .0001 | 0.03 ± 0.019<br>*p* = 0.0705 | 0 |
| **Aci** | 0 | 0.09 ± 0.030<br>*p* = 0.0024 | 0 | 0 | 0 | 0.03 ± 0.011<br>*p* = 0.0163 | 0 | 0.33 ± 0.069<br>*p* < .0001 | 0 | 0.55 ± 0.068<br>*p* < .0001 | 0.12 ± 0.029<br>*p* < .0001 | 0 |
| **All** | 0 | 0 | 0 | 0 | 0 | 0.28 ± 0.069<br>*p* < .0001 | 0 | 0.52 ± 0.057<br>*p* < .0001 | 0 | 0 | 0.18 ± 0.032<br>*p* < .0001 | 0 |
| **Ato** | 0 | 0.13 ± 0.038<br>*p* = 0.0010 | 0 | 0.18 ± 0.49<br>*p* = 0.0002 | 0 | 0.04 ± 0.014<br>*p* = 0.0118 | 0.22 ± 0.052<br>*p* < .0001 | 0.07 ± 0.022<br>*p* = 0.0024 | 0.96 ± 0.044<br>*p* < .0001 | 0.74 ± 0.037<br>*p* < .0001 | -0.30 ± 0.050<br>*p* < .0001 | 0 |
| **Chr** | 0 | 0.11 ± 0.035<br>*p* = 0.0014 | 0 | 0.11 ± 0.0002 | 0 | 0.04 ± 0.012<br>*p* = 0.0131 | 0 | 0.06 ± 0.020<br>*p* = 0.003 | 0 | 0.65 ± 0.044<br>*p* < .0001 | -0.23 ± 0.057<br>*p* < .0001 | 0 |
| **Jeo** | 0 | 0.07 ± 0.025<br>*p* = 0.0040 | 0 | 0 | 0 | 0.98 ± 0.230<br>*p* < .0001 | 0 | 0.44 ± 0.067<br>*p* < .0001 | 0 | 0.43 ± 0.071<br>*p* < .0001 | 0.15 ± 0.032<br>*p* < .0001 | 0 |
| **Lac** | 0 | 0 | 0 | 0 | 0 | 0 | 0 | 0.41 ± 0.065<br>*p* < .0001 | 0 | 0 | 0.14 ± 0.03<br>*p* < .0001 |  |
| **Man** | -0.29 ± 0.072<br>*p* < .0001 | -0.02 ± 0.010<br>*p* = 0.016 | 0 | -0.02 ± 0.011<br>*p* = 0.0386 | -0.08 ± 0.030<br>*p* = 0.0063 | -0.084 ± 0.035<br>*p* = 0.0162 | -0.12 ± 0.037<br>*p* = 0.0014 | -0.04 ± 0.016<br>*p* = 0.0048 | 0 | -0.13 ± 0.039<br>*p* = 0.0005 | (-0.01 ± 0.006)<br>*p* = 0.1017 | 0 |
| **Myc** | 0 | -0.03 ± 0.015<br>*p* = 0.0238 | 0 | -0.45 ± 0.022<br>*p* = 0.0395 | 0 | -0.01 ± 0.005<br>*p* = 0.05 | -0.24 ± 0.075<br>*p* = 0.0016 | -0.02 ± 0.016<br>*p* = 0.0303 | 0 | -0.21 ± 0.067<br>*p* = 0.002 | 0.01 ± 0.006<br>*p* = 0.379 | 0 |
| **Pas** | 0 | -0.03 ± 0.014<br>*p* = 0.02 | -0.25 ± 0.076<br>*p* = 0.001 | -0.45 ± 0.019<br>*p* = 0.0144 | 0 | -0.01 ± 0.004<br>*p* = 0.0451 | -0.06 ± 0.021<br>*p* = 0.001 | -0.02 ± 0.007<br>*p* = 0.0263 | -0.24 ± 0.074<br>*p* = 0.0011 | -0.19 ± 0.058<br>*p* = 0.0012 | -0.20 ± 0.075<br>*p* = 0.0092 | 0 |
| **Pha** | 0 | 0.10 ± 0.031<br>*p* = 0.0017 | 0 | 0.04 ± 0.023<br>*p* = 0.0609 | 0 | 0.03 ± 0.011<br>*p* = 0.0141 | 0.22 ± 0.086<br>*p* = 0.001 | 0.05 ± 0.018<br>*p* = 0.0035 | 0<br>1 | 0.59 ± 0.055<br>*p* < .0001 | -0.17 ± 0.064<br>*p* = 0.0063 | 0 |
| **Pse** | 0 | 0.15 ± 0.045<br>*p* = 0.0011 | 0 | 0.19 ± 0.071<br>*p* = 0.0064 | 0 | 0.04 ± 0.016<br>*p* = 0.0141 | 0 | 0.08 ± 0.025<br>*p* = 0.0024 | 0 | 0.87 ± 0.05<br>*p* < .0001 | -0.02 ± 0.023<br>*p* = 0.3606 | 0 |
| **Psy** | 0 | 0 | 0 | 0 | 0 | 0 | 0 | 0 | 0 | 0 | 0.35 ± 0.048<br>*p* < .0001 | 0 |
| **Rik** | 0 | 0.13 ± 0.040<br>*p* = 0.0009 | 0 | 0.19 ± 0.051<br>*p* = 0.0002 | 0 | 0.04 ± 0.015<br>*p* = 0.0114 | 0.23 ± 0.054<br>*p* < .0001 | 0.07 ± 0.022<br>*p* = 0.0021 | 0 | 0.78 ± 0.034<br>*p* < .0001 | -0.02 ± 0.019<br>*p* = 0.2266 | 0 |
| **Rum** | 0 | 0.17 ± 0.051<br>*p* = 0.0001 | 0 | 0 | 0 | 0.05 ± 0.019<br>*p* = 0.0117 | 0 | 0.09 ± 0.029<br>*p* = 0.0021 | 0 | 0 | 0.03 ± 0.011<br>*p* = 0.0073 | 0 |

**Supplementary table S2.** Continue (Model 3, MP group).
|  | **At** | **Ma** | **Pa** | **Ps** | **Ac** | **Al** | **Ch** | **Day** | **Hi** | **Ph** | **Ps** | **Ri** | **Ru** |
| --- | --- | --- | --- | --- | --- | --- | --- | --- | --- | --- | --- | --- | --- |
| **Ac** | 0.75 ± 0.080<br>*p* < .0001 | 0 | 0 | 0 | 0 | 0 | 0 | 0.29 ± 0.063<br>*p* < .0001 | 0 | 0 | 0 | 0 | 0.6251 ± 0.068<br>*p* < .0001 |
| **At** | 0 | 0 | 0 | 0 | 0 | 0 | 0 | -0.36 ± 0.051<br>*p* < .0001 | 0 | 0 | 0 | 0 | 0.83 ± 0.049<br>*p* < .0001 |
| **Je** | 0.10 ± 0.037<br>*p* = 0.0068 | 0 | 0 | 0.39 ± 0.074<br>*p* < .0001 | 0 | 0 | 0 | 0.16 ± 0.039<br>*p* < .0001 | 0 | 0 | 0 | 0 | 0.50 ± 0.060<br>*p* < .0001 |
| **La** | 0 |  | -0.21 ± 0.073<br>*p* = 0.0041 | 0 | 0.18 ± 0.079<br>*p* = 0.0208 | 0 | 0.20 ± 0.079<br>*p* = 0.0132 | 0.045 ± 0.023<br>*p* = 0.0471 | 0 | 0 | 0 | 0 | 0 |
| **Ma** | 0 | 0 | 0 | 0 | 0 | 0 | 0 | -0.19 ± 0.077<br>*p* = 0.0165 | 0 | 0 | 0 | 0 | 0 |
| **My** | -0.48 ± 0.055<br>*p* < .0001 | -0.16 ± 0.057<br>*p* = 0.0053 | 0 | 0 | 0 | 0 | 0 | 0.02 ± 0.035<br>*p* < .0001 | 0 | 0 | 0 | -0.35 ± 0.058<br>*p* < .0001 | -0.40 ± 0.053<br>*p* < .0001 |
| **Pa** | 0 | 0 | 0 | 0 | 0 | 0 | 0 | -0.22 ± 0.076<br>*p* = 0.005 | 0 | 0 | 0 | 0 | 0 |
| **Ps** | 0.26 ± 0.081<br>*p* = 0.0014 | 0 | 0 | 0 | 0 | 0 | 0 | 0.42 ± 0.055<br>*p* < .0001 | 0 | 0 | 0 | 0 | 0.60 ± 0.051<br>*p* < .0001 |

**Supplementary Table S3.** Standardized total effects from *Lactobacillus* to the abundances of other bacteria.The standardized total effects of Lactobacillus are shown in Table S2 for Model 1 (BT group) and Model 2 (CTRL group). *Lactobacillus* was not a predictor variable in Model 3 (MP group).

|  | Model 1 (BT) | Model 2 (CTRL) |
| --- | --- | --- |
| *Acetitomaculum* | 0.0237 ± 0.008837<br>*p* = 0.007316 | 0.2949 ± 0.1018<br>*p* = 0.003762 |
| *Acinetobacter* | 0.2379 ± 0.0615<br>*p* = 0.000108 | 0.0257 ± 0.0107<br>*p* = 0.0166 |
| *Alloprevotella* | 0.025 ± 0.009363<br>*p* = 0.007598 | 0.2794 ± 0.071<br>*p* <.0001 |
| *Atopostipes* | 0.0305 ± 0.0112,<br>*p* = 0.006419 | 0.0348 ± 0.0138<br>*p* = 0.0116 |
| *Christensenellaceae_R7_group* | 0.0946 ± 0.0251<br>*p* 0.00016 | 0.0307 ± 0.0124<br>*p* = 0.0136 |
| *Jeotgalibaca* | 0.0119 ± 0.0196<br>*p* = 0.5419 | 0.9835 ± 0.2532<br>*p* = 0.000102 |
| *Mannheimia* | 0.1875 ± 0.0758<br>*p* = 0.0134 | -0.0844 ± 0.0367<br>*p* = 0.0216 |
| *Mycoplasma* | 0 | -0.00966 ± 0.004982<br>*p* = 0.0525 |
| *Pasteurella* | 0 | -0.0883 ± 0.004454<br>*p* = 0.0474 |
| *Phascolarctobacterium* | 0.0688 ± 0.0258<br>*p* = 0.007703 | 0.0275 ± 0.0113<br>*p* = 0.0149 |
| *Pseudomonas* | 0 | 0.0409 ± 0.0163<br>*p* = 0.0123 |
| *Psychrobacter* | 0.0349 ± 0.0128<br>*p* = 0.006275 | 0 |
| *Rikenellaceae_RC9_gut_group* | 0.0833 ± 0.0225<br>*p* = 0.00021 | 0.0366 ± 0.0145<br>*p* = 0.0116 |
| *Ruminococcaceae_UCG005* | 0.1251 ± 0.0352<br>*p* = 0.00038 | 0.047 ± 0.0186<br>*p* = 0.0115 |

**Supplement Fig. S1.**
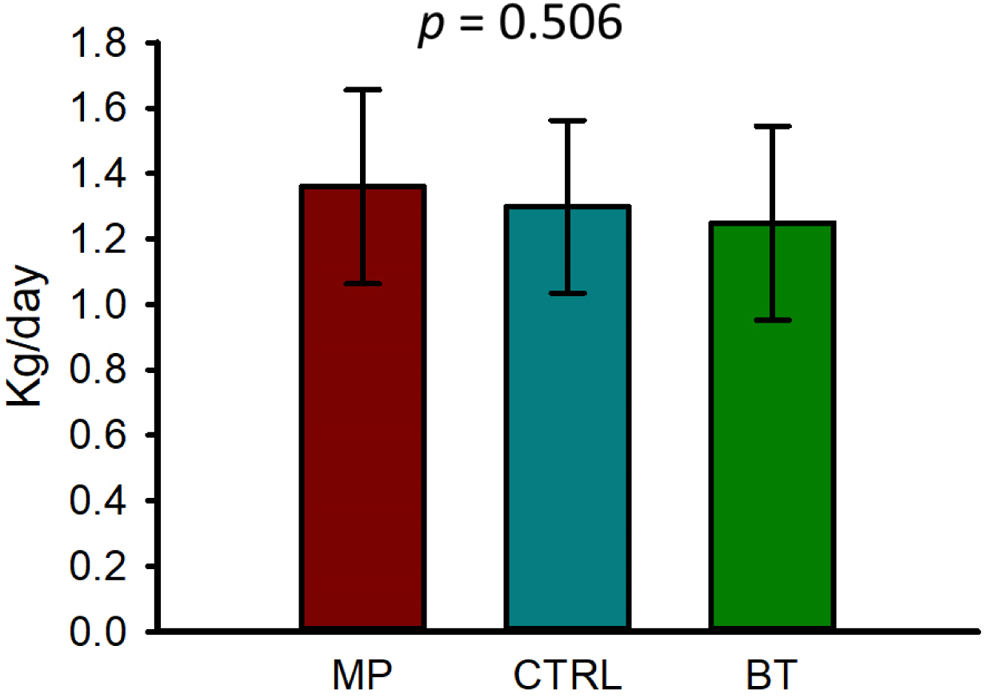
Average daily gain of the cattle over 42 days. On day 1, cattle were treated with intranasal bacterial therapeutics (BT), intranasal PBS (CTRL), or subcutaneous tulathromycin (MP) (n = 20 per group). The bar plot represents the mean average daily gain. Error bars indicate ± standard error of the mean.

**Supplement Fig. S2.**
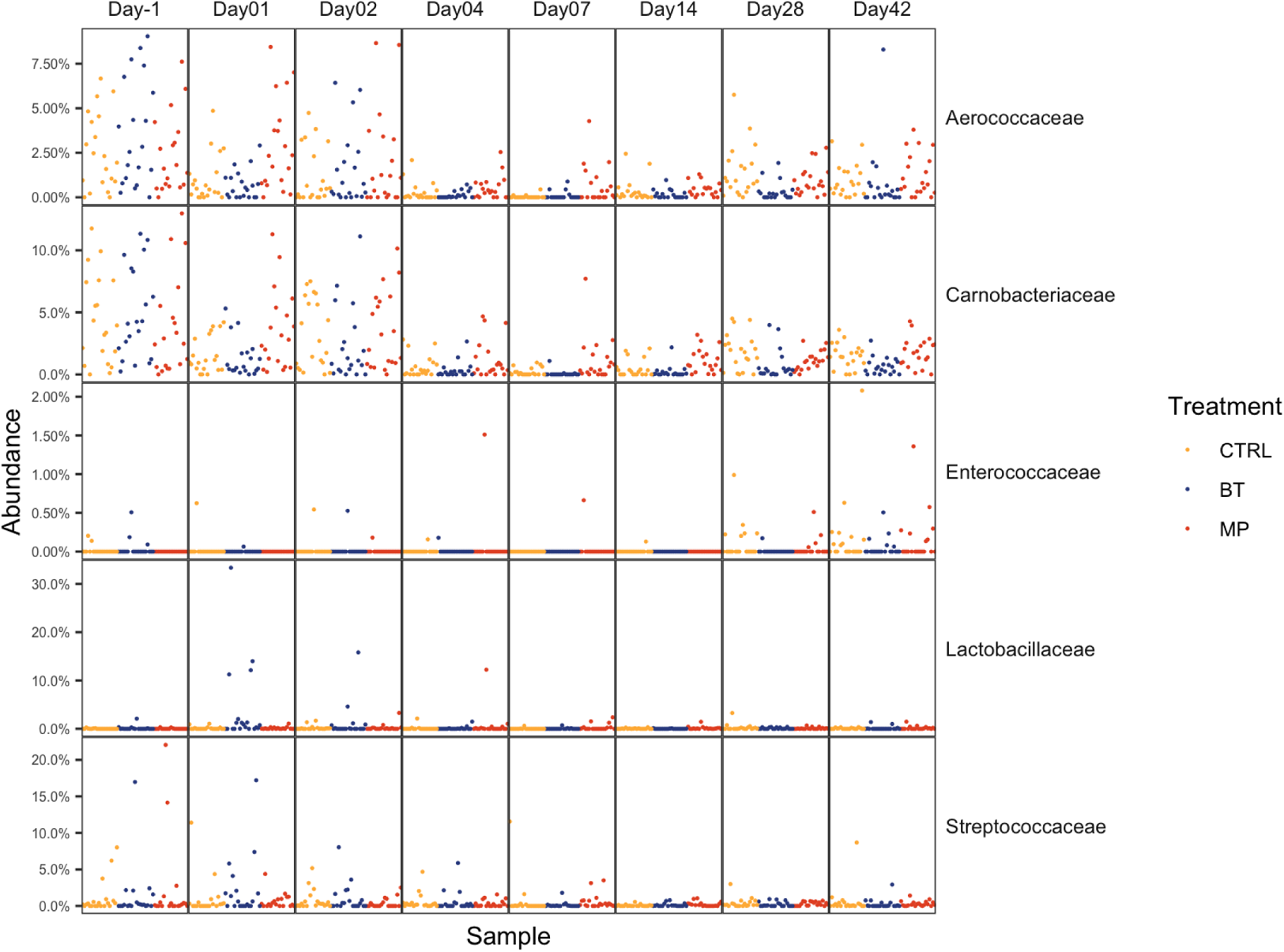
The relative abundance of families within the order *Lactobacillales* in the nasopharyngeal of cattle. On day 1, cattle were treated with intranasal bacterial therapeutics (BT), intranasal PBS (CTRL), or subcutaneous tulathromycin (MP) (n = 20 per group).

**Supplement Fig. S3.**
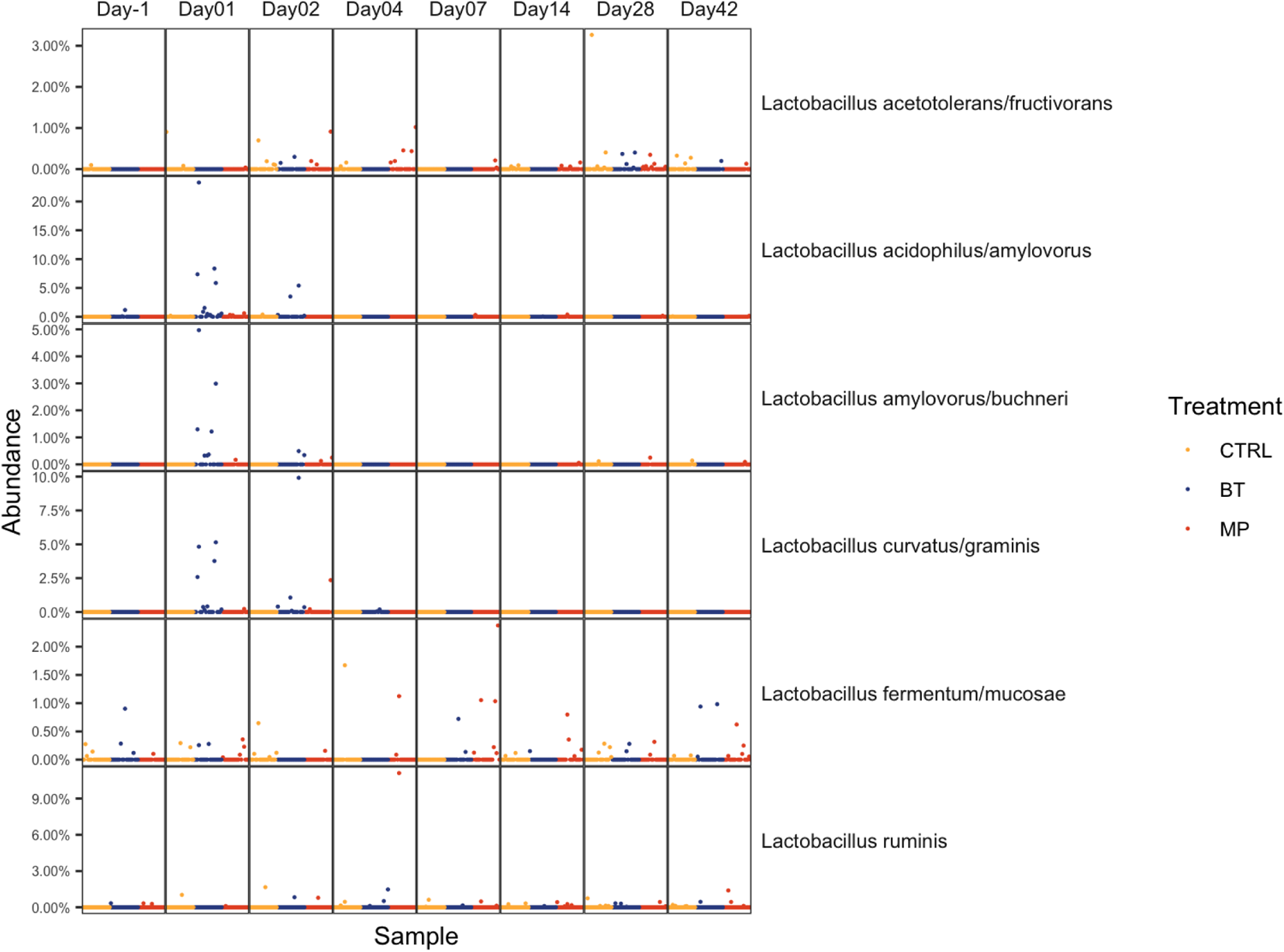
The relative abundance of different *Lactobacillus* species in the nasopharynx of cattle, determined by 16S rRNA sequencing. On day 1, cattle were treated with intranasal bacterial therapeutics (BT), intranasal PBS (CTRL), or subcutaneous tulathromycin (MP) (n = 20 per group).

**Supplement Fig. S4.**
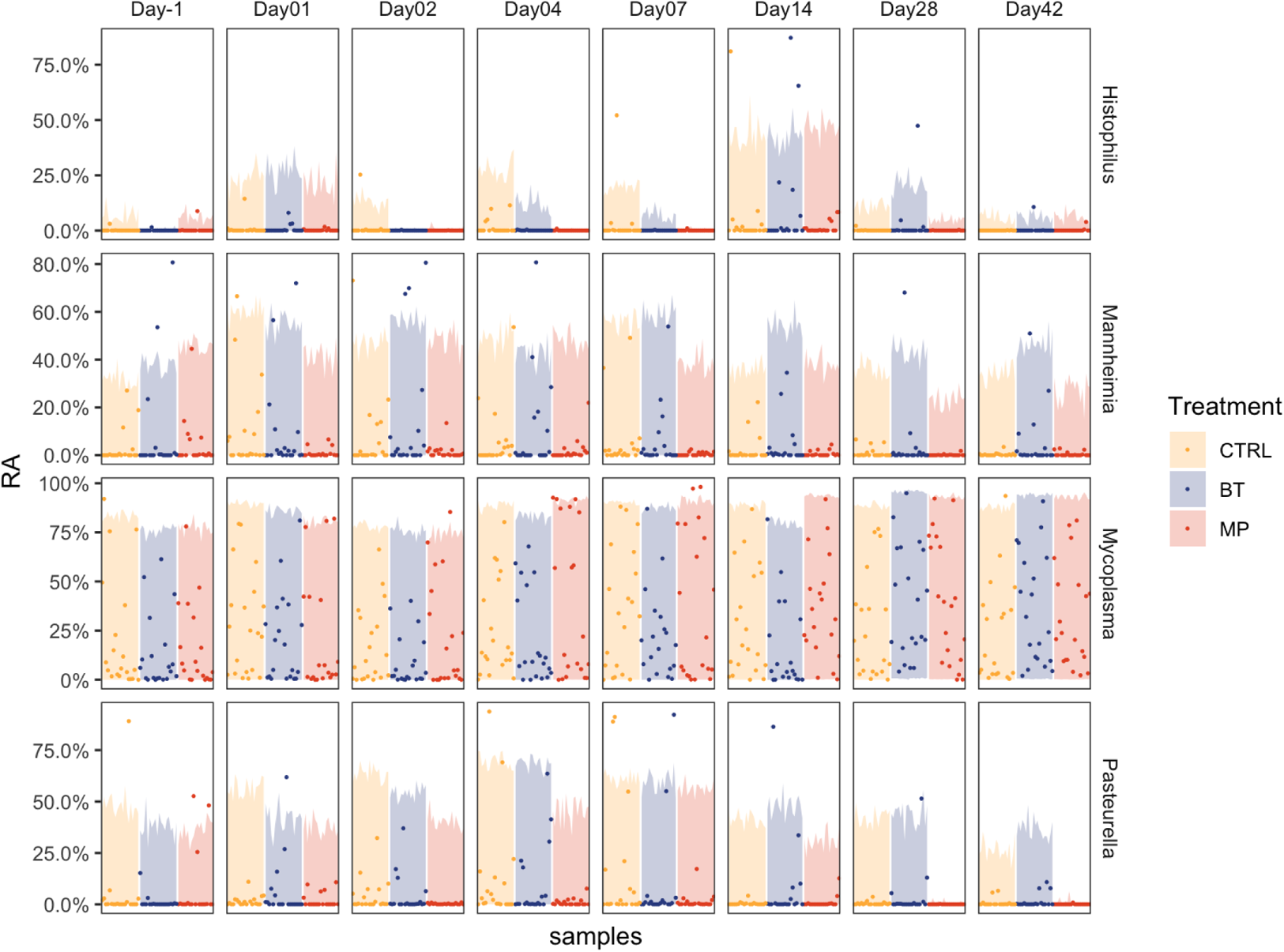
Relative abundance of genera associated with bovine respiratory disease in the nasopharynx of cattle, determined by 16S rRNA gene sequencing. On day 1, cattle were treated with intranasal bacterial therapeutics (BT), intranasal PBS (CTRL), or subcutaneous tulathromycin (MP) (n = 20 per group).

**Supplement Fig.S5.**
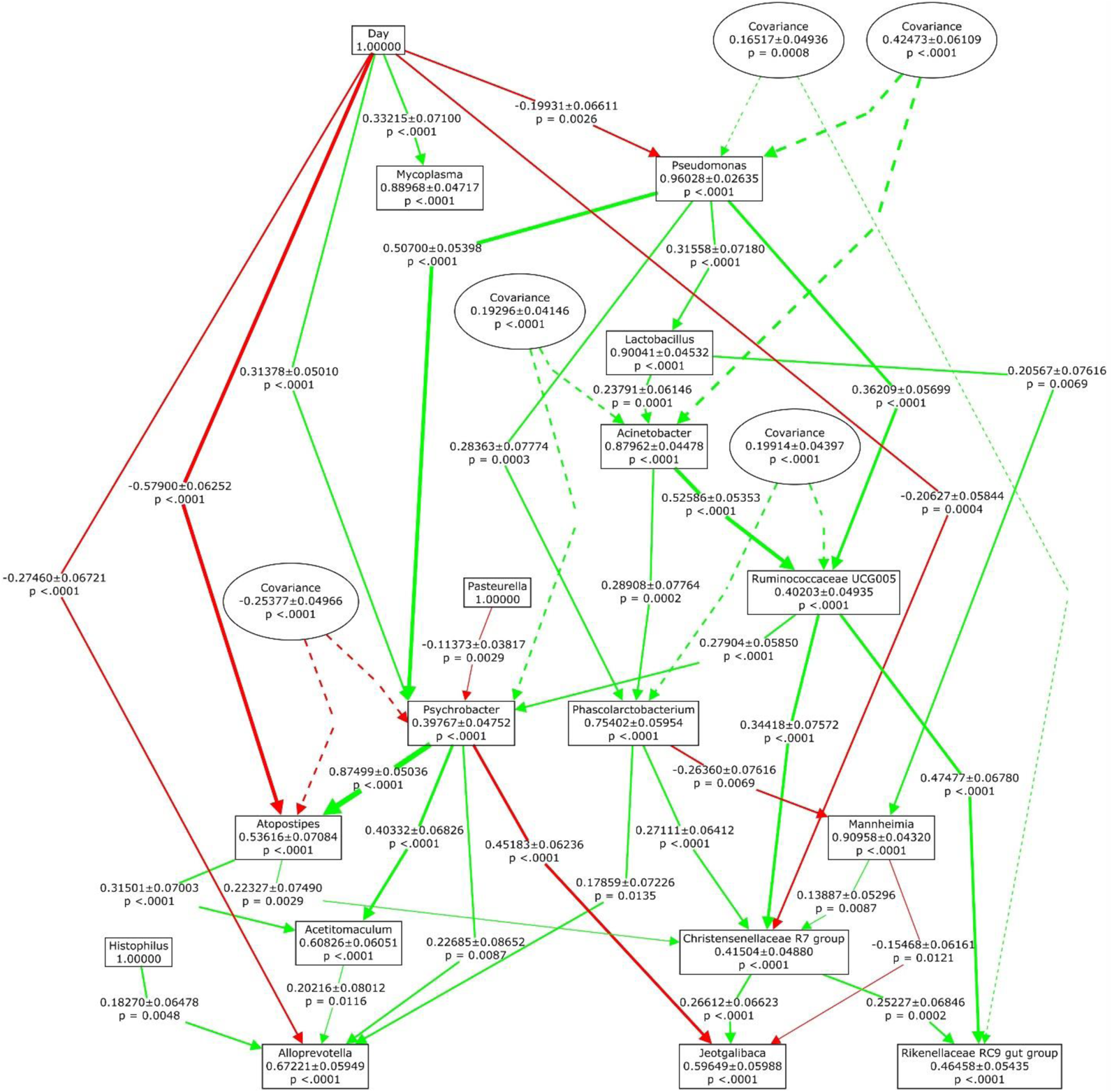
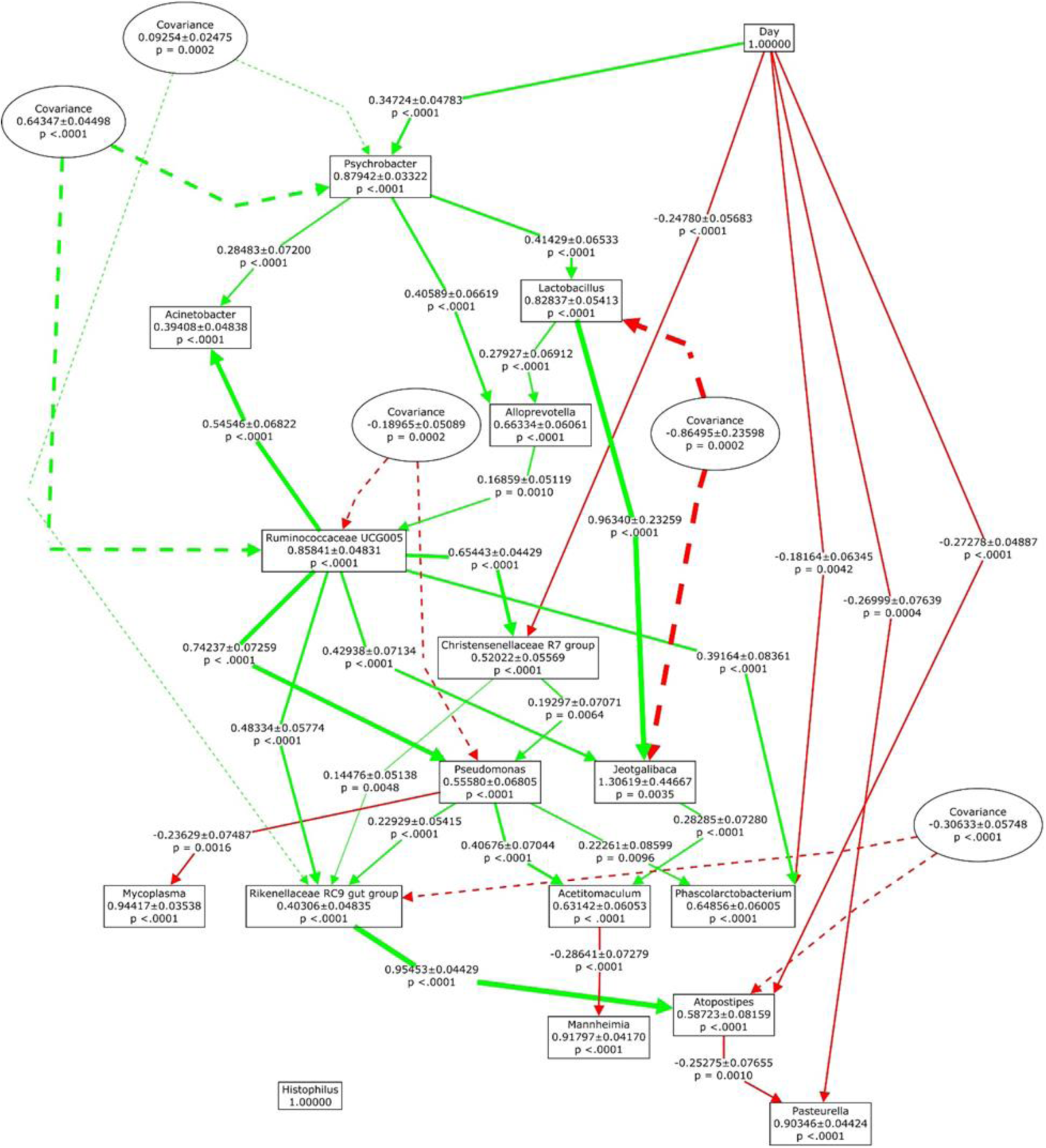
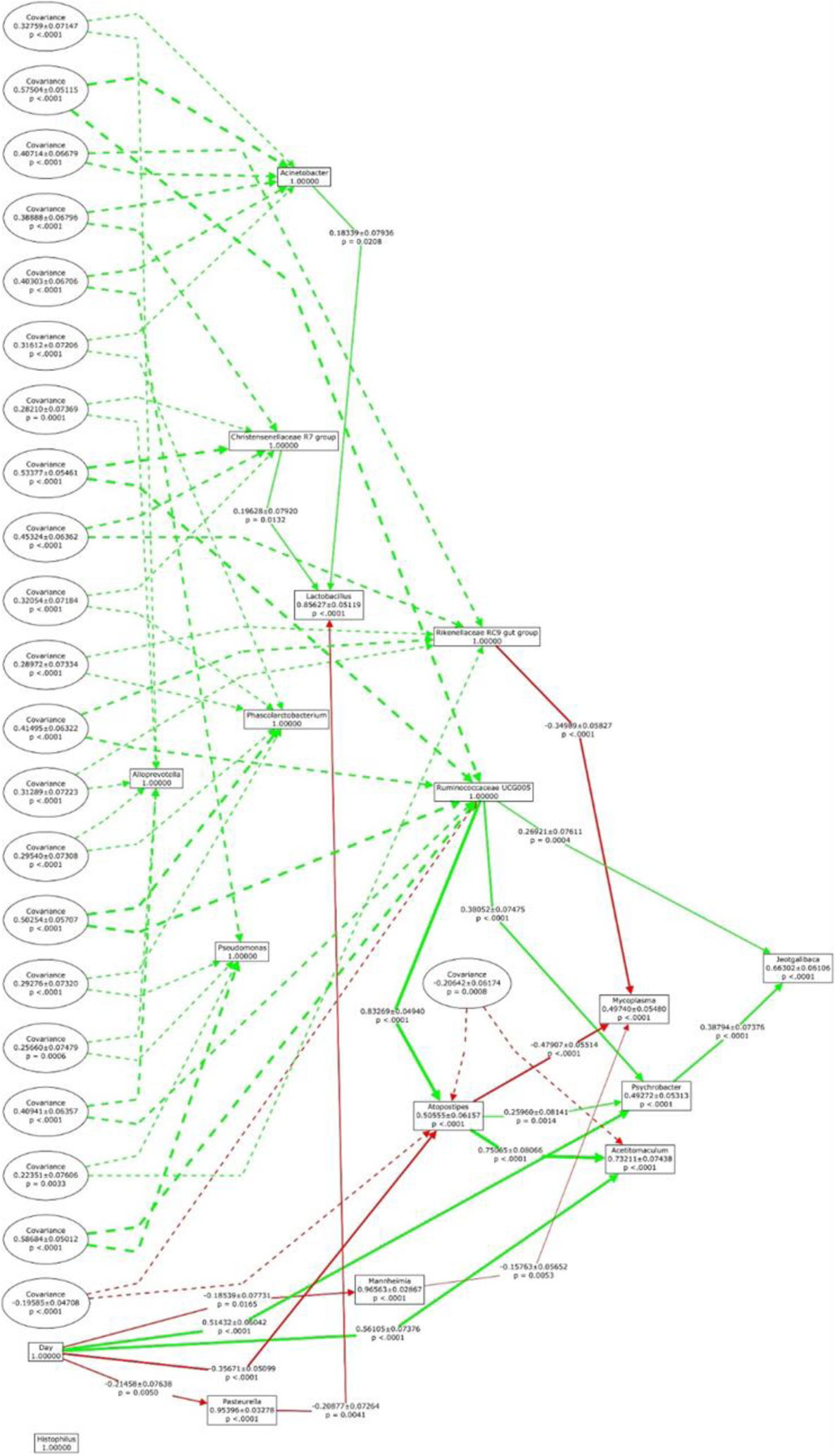
Path diagram of models showing the recursive structure of causal relationships among 16 selected genera in the nasopharyngeal microbiota of calves that received an intranasal inoculation of either bacterial therapeutics (BT) (**A**) or PBS (CTRL) (**B**), or subcutaneous metaphylaxis (MP) (**C**) (n = 20 per group). Variances of measured variables (relative abundances of genera and time) with standard errors and P values are shown in squares. Causal relationships that are implied by the model are shown as solid lines with arrows that indicate the direction of causation. Causal paths are labelled with the standardized path coefficients, standard errors, and P values. Covariance terms are shown in ovals with dashed arrows between the variables that co-vary. Green line represents the positive effect, whereas the red line represents the negative effect. The thickness of the solid line represents the strength of the effect. The genera selected for analysis were based on the 15 most relatively abundant genera observed from 16S rRNA sequencing plus *Lactobacillus* genus.

